# A spatial gene expression signature of the mouse brain post-injury at the focal point of contusion

**DOI:** 10.1101/2025.07.21.665943

**Authors:** S. K. Kounelis-Wuillaume, A. M. Frank, E. Goguet, C. Alba, G. Sukumar, M. D. Wilkerson, C. L. Dalgard, J. T. McCabe, M. L. Doughty

## Abstract

Traumatic brain injury (TBI) results from a primary injury that impacts the brain in a spatially-dependent manner. Here we investigated the topographical relationship of early transcriptional responses to a single, focal TBI in mice by controlled cortical impact (CCI). Guided by the presence of the anterior commissure (AC) in coronal sections at the rostro-caudal point of impact, we compared gene expression changes in the neocortex (CTX) and corpus callosum-external capsule (CC-EC), striatum (STR) and AC. Injury-induced gene expression changes were detected in the CTX, CC-EC and STR but not AC and were principally segregated based on cytoarchitecture, and secondarily by proximity to the site of impact. Additionally, unbiased spatial clustering revealed a positive relationship between proximity to the impact and the number of acutely differentially expressed genes within the laminar CTX. Gene pathways for interferon gamma response and for leukocyte-mediated migration and immunity were acutely enhanced across the injured CTX, CC-EC and STR. Within 1-week post-injury, transcriptional responses to injury in the CTX and CC-EC included gene pathways for adaptive T- and B-cell mediated immunity, whereas gene expression changes in the STR were largely resolved. Next, we examined the effects of systemic depletion of neutrophils and monocytes on spatial gene expression changes in the injured brain. This led to the upregulation of gene pathways functioning in synaptic transmission and an alternating down- and then upregulation of genes functioning in ribosomal messenger RNA translation and aerobic metabolism in mitochondria. These data suggest infiltrating neutrophils and monocytes play an evolving, multifaceted role in modulating the metabolic, transcriptional, and synaptic activity of brain tissue post-injury.

## Introduction

Traumatic brain injury (TBI) is a leading cause of morbidity and mortality in the United States [1] that results in a wide range of pathophysiological outcomes [2]. TBI induces both a primary injury, caused by the external forces acting at the moment of insult, and a subsequent secondary injury initiated by the primary trauma. TBI is recognized as a chronic disease with a broad spectrum of clinical and pathological sequelae whose course is largely dictated by the severity of secondary injury ^[3]^.

The acute phases of secondary injury following experimental TBI are characterized by the activation of neuroinflammatory pathways, the disruption of the blood brain barrier (BBB) with resultant brain tissue edema, and the infiltration of peripheral immune cells (PICs) into damaged brain tissue. PICs infiltrate the injured brain parenchyma across the dysfunctional BBB in response to chemoattractant cues such as damage/disease associated molecular patterns (DAMPs) and cytokines/chemokines (C/Cs) from the damaged resident neurons and neuroglial cells [4].

Neutrophils are the first and most numerous PICs to respond to TBI, entering the injured tissue within minutes to hours post-injury [5, 6]. Neutrophil recruitment can assist in limiting the spread of injury by phagocytosis [7] but has also been demonstrated to be neurotoxic, propagating neuroinflammation ^[8, 9]^ and releasing factors that contribute to BBB breakdown [10, 11]. Neutrophil infiltration peaks approximately 2 days post-injury (DPI) in preclinical TBI rodent models. As neutrophil recruitment declines, monocytes begin to infiltrate the brain and resident microglia and astrocytes accumulate at the site of injury [12, 13]. Infiltrating monocytes adopt macrophage phenotypes in the brain with a spectrum from pro- to anti-inflammatory function [14]. Monocytes are predominantly detrimental to neuronal survival and function in the acute (hours-days) recovery period after TBI, directing a type I interferon (IFN) response in microglia [15], but conversely play a more regenerative role during the chronic (weeks-months) TBI recovery phase [9, 16–18]. Finally, depending on the severity of experimental injury, adaptive immune cells, such as T and B cells, can also penetrate the brain during the first week post-injury with opposing pro- and anti-inflammatory signaling effects amongst T cell subsets [19, 20]. Thus, post-injury infiltration of immune cells can be either detrimental or beneficial to repair after TBI depending on the timing of their infiltration, the cell types involved, and the cells’ specific phenotypes [21, 22]. Additionally, since TBI is dependent upon physical forces interacting with the brain, the heterogeneity of responses at the cellular level to immune cell infiltration are predicted to translate into differences at the spatial level as well. Indeed, advanced neuroimaging supports the hypothesis that spatially distinct alterations in cellular damage, remodeling and repair pathways is a hallmark of clinical TBI [23].

In this study, we tested the hypothesis that attenuating the infiltration of neutrophils and monocytes into the brain post-injury would alter transcriptional responses to trauma in a spatially-dependent manner. To examine the spatial molecular response to injury alone, we first mapped early post-injury transcriptional responses in a well-characterized mouse model of focal contusion injury using spatial transcriptomics. Guided by the presence of the anterior commissure (AC) in coronal sections at the rostro-caudal focal point of impact, we mapped gene expression at the contusion epicenter at 2 and 7 DPI in adult mice. As expected, this approach revealed spatiotemporally distinct transcriptional responses in the cortex, striatum and damaged white matter of the forebrain. Next, we systemically treated mice with antibodies to Granulocyte Receptor-1 (anti-GR-1) to deplete circulating and brain-infiltrating neutrophils and monocytes and repeated our spatial transcriptomics analysis. GR-1^+^ immune cell depletion resulted in the surprising upregulation of genes functioning in synaptic activity in the injured forebrain, along with the acute suppression and subsequent increased activation of gene pathways functioning in ribosomal messenger RNA translation and mitochondrial metabolism in the CTX.

## Methods and Materials

### Animal Care and Surgery

#### Animals

Male C57BL/6J mice (strain# 000664) were obtained from The Jackson Laboratory and allowed to acclimate after arrival for at least 3 days before commencing procedures. Animals were group-housed, kept on a 12:12 hour light: dark schedule, and food and water were available *ad libitum*. Surgical procedures and behavioral testing were performed during the light phase of the diurnal cycle. Animal housing facilities were accredited by the Association for the Advancement and Accreditation of Laboratory Animal Care and all animal procedures described were approved by the Institutional Animal Care and Use Committee (IACUC) at the Uniformed Services University of the Health Sciences (Bethesda, MD) under protocol # APG-23-893.

#### Controlled Cortical Impact

The CCI and Sham control models chosen in this study have been previously described and characterized [24]. Mice 16 to 20 weeks of age were anesthetized with 3% isoflurane (Forane, Baxter Healthcare Corporation, Deerfield, IL). After cessation of corneal and pedal reflexes, the scalp fur was clipped. Mice were then placed into a stereotaxic device with an incisor bar and atraumatic ear bars. Anesthesia (≈2-2.5% isoflurane) was maintained via a flow-through nose cone, and anti-microbial ophthalmic ointment was applied. The scalp was sterilized with betadine and 70% ethanol, following which a midline incision was performed. The skin was pulled back, and fascia cleared to expose the left side of the skull. A 5 mm craniectomy centered at 2.0 mm lateral to bregma (Anterior-Posterior = 0 mm) was cut using a high-speed rotary tool with a 0.6 mm burr drill bit. Cortical impact was performed with the Impact One™ device (Leica Microsystems, Buffalo Grove, IL) using a 3 mm diameter probe at a 15-degree angle relative to the sagittal plane. Impact velocity was set at 1.5 m/s, displacement depth was 1 mm, and dwell time was 100 ms. Sham controls received scalp incision and clearing of the skull fascia under anesthesia of similar time duration to the CCI surgeries. Following CCI or Sham surgery, the incision was sutured with simple interrupted stiches (monocryl, 4-0, Ethicon) without replacement of the bone flap. Topical antibiotic ointment (triple antibiotic ointment: bacitracin zinc, neomycin sulfate, and polymyxin B sulfate, Globe) was applied over the incision site. Mice were then allowed to recover in a pre-warmed cage with softened chow available on the cage floor until fully alert and ambulatory. All mice received acetaminophen in their drinking water (Children’s Mapap, Livonia, MI; 1 mg/ml) beginning from the timepoint of acclimation in the surgical suite until 24 hours after the final surgery.

#### Systemic depletion of GR-1^+^ leukocytes

GR-1 is comprised of two structurally-related glycosylphosphatidylinositol-anchored proteins of the Ly-6 superfamily, Ly6G and Ly6C. Beginning on the day of CCI or Sham surgeries and continuously thereafter until either 2 or 7 DPI endpoint, mice received 250 µg of monoclonal IgG2b rat anti-mouse GR-1 antibody (clone RB6-8C5, BioXCell, cat# BE0075) daily by intraperitoneal (ip) injection [25–27]. Controls received the same regimen and dose of ip injections of an isotype monoclonal IgG2b rat anti-keyhole limpet hemocyanin antibody (referred to from herein as Control antibody; clone LTF-2, BioXCell, cat# BE0090). Limpet hemocyanin is not expressed in mammals and the antibody is therefore non-reactive in mice. Mice receiving anti-GR-1 or Control antibody treatment were weighed daily beginning just prior to ip injection until the day before reaching their study endpoint. Mice body weights were compared between each group over time via a Mixed Effects ANOVA in Graphpad Prism 10.1.0. p values ≤ 0.05 were considered significant.

### Flow Cytometry

#### Flow Cytometry: Isolation of Circulating Leukocytes

Blood was collected directly into ethylenediaminetetraacetic acid (EDTA)-coated tubes (BD Microtainer® K_2_EDTA tubes, BD Biosciences) during rapid decapitation. The blood from two mice in an experimental group at each timepoint were pooled into a 50 mL centrifuge tube and incubated in either i) 5 mL 1X Red Blood Cell (RBC) Lysis Buffer prepared fresh from 10X RBC Lysis Buffer (Cell Signaling Technology) or ii) 10 mL of 1X Ammonium-Chloride-Potassium (ACK) Lysis buffer (Quality Biological) for 15 min at room temperature (RT), protected from light. At the end of incubation, 20 mL of 1X PBS was added per tube to slow the reaction. Cells were then spun down in a pre-cooled swing-bucket Eppendorf 5810R Centrifuge at 4°C and 500 x g for 5 min. The supernatant was removed and cells were then washed in 5 mL ice-cold 1X Phosphate Buffered Saline (PBS) and spun down at 500 x g for 5 min at 4°C. Blood leukocytes were then resuspended in 2 mL 1X PBS and placed on ice until cell counting.

#### Flow Cytometry: Isolation of Splenocytes

Following euthanasia and collection of blood and/or brain tissues, spleens were isolated and kept in a 1.5 mL centrifuge tube filled with 1X PBS on ice until splenocyte isolation could be performed. Two spleens per group were combined for processing following this step. Spleens and buffer were then transferred onto a 70 µm-cell strainer (Midwest Scientific) and homogenized on top of the filter with the plunger of a 3 mL syringe. The cell strainer was then rinsed with 10 mL of 1X PBS and cells were pelleted by centrifugation at 500 x g for 5 min at 4°C. The supernatant was removed and cells were resuspended in either: i) 10 mL of freshly prepared 1X RBC lysis buffer, or ii) 6 mL 1X ACK lysis buffer and incubated for 15 min at RT protected from light. At the end of incubation, 10 mL of 1X PBS was added per tube to slow the reaction. Cells were then spun down at 500 x g for 5 min at 4°C. The supernatant was removed and cells were then washed in 5 mL ice-chilled 1X PBS and spun down at 500 x g for 5 min at 4°C. Splenocytes were then resuspended in 10 mL 1X PBS and placed on ice until cell counting.

#### Preparation of Single Cell Suspensions of Isolated Brain Tissue for Flow Cytometry

Isolated brain tissue was processed into single cell populations for flow cytometry using a modified version of the protocol published by Liu et al. [28] and recommendations from the MACS Adult Brain Dissociation Kit Protocol (Miltenyi Biotech, cat# 130-107-677). All plasticware, including pipet tips, was always initially rinsed with coating buffer comprised of 3% w/v bovine serum albumin (BSA) in 1X Dulbecco’s Phosphate Buffered Saline – Calcium and Magnesium Free (DPBS-CMF, Thermo Fisher). Mice were anesthetized under isoflurane (4%) and transcardially perfused with at least 20 mL of ice-cold perfusion solution made of 30 mL of DPBS-CMF plus 2 UI/mL of heparin (Fresenius Kabi USA, LLC). Brains were then rapidly dissected from the skull and cut along the mid-sagittal suture into an ipsilateral (left, contused or sham) and contralateral hemisphere with a sterile scalpel blade. The ipsilateral brain hemisphere was stored separately from contralateral hemispheres in C-tubes filled with ice-cold and freshly prepared Enzyme Mix 1 comprised of 1.9 mL Buffer Z and 50 µL Enzyme P from the MACS Adult Brain Dissociation Kit.

Once all brain tissue was isolated, brain hemispheres were gently minced with sterile scissors one-by-one on ice in their C tube. Enzyme Mix 2, comprised of 20 µL Buffer Y and 10 µL Enzyme A per brain hemisphere, was added to each sample on ice which were immediately placed onto the gentleMACS^TM^ OctoDissociator after tapping all of the tissue and liquid to the bottom of the inverted tube. The tissue was then run on the 37C_ABDK_01 gentleMACS^TM^ OctoDissociator program which is optimized for brain tissue > 100mg. During the program, sterile 70 µm-cell strainers were placed onto 50 mL tubes in ice and pre-wet with 500 µL ice-cold DPBS-CMF. Following program completion, samples were immediately removed from the OctoDissociator and placed onto ice. 9.5 mL of ice-cold DPBS-CMF was then added to each C tube. The C tube was inverted several times before adding the sample onto a cell strainer for collection of the flow-through in the 50 mL tube. The strainers were disposed of and the 50 mL tubes were spun down at 300 x g for 10 min at 4°C. The supernatant was aspirated from each sample slowly and without disturbing the cell pellet. Next, the Debris Removal protocol and Myelin Debris Removal protocols reported in Liu et al. 2021 were performed exactly as detailed in ^32^ to obtain a single-cell suspension. Each brain hemisphere sample was run on its own LS Column. However, all samples in a group were collected into the same 15 mL conical tube such that Sham ipsilateral or contralateral hemispheres had n=2 and CCI ipsilateral or contralateral hemispheres had n=4 per tube. Samples were then spun down for 10 min at 300 x g at 4°C and resuspended in 600 µL (n=2) or 1.2 mL (n=4) loading buffer comprised of 0.04% (w/v) BSA in 1X DPBS-CMF. Samples were kept on ice until proceeding with cell counting and flow cytometry.

#### Cell Counts

Cells were counted on a QuadCount (Accuris, Edison, NJ, USA) using the preset for peripheral blood mononuclear cells. 10 µL of each cell suspension was combined with 10 µL of Trypan Blue (Invitrogen, Carlsbad, CA, USA) and 20 µL was loaded into the chamber of a QuadCount slide immediately prior to counting.

#### Flow Cytometry: Identification of Microglia and Brain-Infiltrating Leukocytes

All centrifugation steps were performed at 4°C on an Allegra® X-15R centrifuge (Beckman Coulter, Brea, CA, USA) and cells were incubated on ice and protected from light. All buffers were chilled on ice during use. 10^6^ cells from each experimental group were transferred to individual 5 ml polypropylene-flow tubes. If 10^6^ cells were not available for an experimental group, the maximum number of cells were transferred and were at least 500,000. To create unstained controls, viability controls, single color controls (SCCs), and fluorescence minus one (FMO) controls, cells from each experimental group were mixed together and 10^6^ cells of this mixture were added per tube. Each flow tube was then washed once in 1 mL of 1X PBS and spun down at 400 x g for 5 min. Supernatant was removed and cells were resuspended in 1 mL 1X PBS. For viability controls, 500 µL of each cell suspension was removed into a 1.5 mL centrifuge tube and placed in a heat block at 80°C for 15 min (blood, spleen samples) or up to 30 min (brain leukocyte samples). After heat-shock, tubes were incubated on ice for 1 min before being added back to their respective viability control flow tubes.

Next, 10 µL of LIVE/DEAD Fixable Blue (Thermo Fisher) master mix comprised of 0.5 µL LIVE/DEAD Fixable Blue combined with 9.5 µL 1X PBS was added to all flow tubes except the unstained controls and the SCCs. Cells were incubated on ice for 30 min after which all tubes were washed with 1 mL of 1X PBS and spun down at 400 x g for 5 min. During this time, bead SCCs were created by adding two drops of UltraComp eBeads (Invitrogen, Waltham, MA, USA) to 100 µL 1X PBS + 0.5% BSA. An Fc Block master mix was also prepared by combining 49 µL 1X PBS/0.5% BSA with 1 µL Mouse TruStain FcX (clone 2.4G2; BD Biosciences, San Jose, CA, USA) per flow tube. After the wash, all cells were resuspended in 50 µL of Fc Block Master Mix and incubated for 10 min, except the SCCs, unstained, and viability controls which were resuspended in 100 µL 1X PBS/0.5% BSA and kept on ice.

During this time, a master mix of our flow cytometry panel antibodies was created. To confirm depletion of both circulating and brain-infiltrating GR-1^+^ leukocytes, we developed a flow cytometry panel with the help of Cytek Biosciences (Fremont, CA, USA), see **Supplementary Materials Table A**. The chosen concentration of antibody was determined by titration of at least 3 dilutions on a mixture of our experimental splenocytes. The concentration used in our flow cytometry panel had the highest stain index (SI). SI was calculated as the mean fluorescent intensity (MFI) of the positive cell population minus the MFI of the negative cell population divided by 2 times the standard deviation of the negative cell population.

Following this incubation, all experimental tubes received flow staining master mix and were incubated for 45 min on ice and protected from light. During this time, all SCCs and FMO controls received appropriate amounts of antibodies. Following incubation, cells were rinsed with 1 mL 1X PBS/0.5% BSA at 400 x g for 5 min then resuspended in 300 µL 1X PBS/0.5% BSA. All samples were measured in the Uniformed Services University Biomedical Instrumentation Center Flow Cytometry Resource Center (Bethesda, MD, USA) on the Cytek Aurora (Cytek Biosciences). Each experiment underwent unmixing and correction for autofluorescence before being exported as an FCS file for further analysis. Quantification of cell populations was performed using the FlowJo software (FlowJo LLC, BD Life Sciences, Ashland, OR, USA).

### Visium Spatial Gene Expression Analysis

#### Tissue Preparation for Spatial Transcriptomics

Mice were deeply anesthetized with inhaled 4% isoflurane in O_2_ vehicle gas. When mechanical stimuli failed to produce paw withdrawal or eye blink reflexes, mice were rapidly decapitated, and the brain was quickly removed from the skull. All tissue preparation was performed following *Visium Spatial Protocols – Tissue Preparation Guide* (Document number CG000240, Rev C, 10X Genomics). In summary, once the brain was isolated, excess blood was removed from the tissue by light dabbing with a Kimwipe. The brain was then covered in a layer of pre-chilled Tissue-Tek® optimum cutting temperature (OCT) compound (Sakura) and then placed into a pre-chilled tissue cassette already holding a layer of OCT before being completely covered by pre-chilled OCT. The tissue cassette was then placed into a cooled isopentane (2-methylbutane, Millipore Sigma) bath surrounded by liquid nitrogen and dry ice to rapidly freeze the tissue. Tissue was then stored for at least 24 hours at −80°C before being sectioned on a Leica Cryostat (Leica, Germany) with the cryostat temperature set to −20°C and the tissue block temperature set to −15°C. All materials and tissues were removed from −80°C storage onto dry ice and then immediately placed in the cryostat and allowed to acclimate for at least 30 min before sectioning. Brain tissue was sectioned in the coronal plane. See **Supplementary Materials Table B** for Visium experimental group sizes.

#### RNA quality control

For determination of RNA quality, 10 sections of 20 µm thickness were collected into 1.5 mL centrifuge tubes and stored at −80°C until isolation was performed. For placement onto Visium Fresh Frozen Spatial GEX slides (PN-2000233) or Visium Fresh Frozen Spatial Tissue Optimization Slides (PN-3000394), brains were scored along the midline and the left hemisphere was sectioned at 12 µm thickness and placed within the 6.5 x 6.5 mm frames on the GEX or Optimization slides at the point the AC was visible (approximately bregma 0.0 mm and anterior-posterior 0.0 mm). This ensures capture of the epicenter of the impact in our CCI mice. Both CCI and Sham brain tissues were placed on each Visium GEX slide while only CCI tissue was used on Visium Optimization slides.

#### Hematoxylin & Eosin (H&E) Staining and Imaging

Histology and Imaging was performed according to the *Methanol Fixation, H&E Staining & Imaging for Visium Spatial Protocols for Fresh Frozen tissue* (Document number CG000160, Rev B, 10X Genomics). Following sectioning, the slides were placed on a 37°C pre-warmed adaptor provided by 10X Genomics fit inside a T100 thermal cycler (BioRad, Hercules, CA, USA) for 1 min. Tissues were then immediately fixed in pre-chilled methanol at −20°C for 30 min. After fixation, slides were removed from the methanol to room temperature and excess liquid removed with a Kimwipe before applying 500 µL isopropanol (Millipore Sigma) for 1 min. The isopropanol was carefully poured off the slides and tissues were allowed to dry for ≈ 5-6 min. H&E staining was then performed as stated in the 10X Genomics protocol using Mayer’s Hematoxylin (Electron Microscopy Sciences) and Eosin Y (Electron Microscopy Sciences). For the bluing step, DEPC-treated and autoclaved tap water was used. Washes performed between staining steps used MilliQ water. Following all staining steps and washes, slides were dried for ≈ 5 min prior to imaging under brightfield at 20X without a coverslip on an Axioscan (Zeiss, Germany).

#### Visium Fresh Frozen (FF) Spatial Tissue Optimization

Optimization was conducted according to the *Visium Spatial GEX Tissue Optimization User Guide* (Document number CG000238, Rev D, 10X Genomics). The contused hemispheres of CCI mouse brain tissue at 2 and 7 DPI were used on two separate optimization slides for determination of the best permeabilization enzyme incubation time. Following tissue sectioning, H&E staining as described above, and imaging of the optimization slide, tissues were permeabilized for various times as described in the protocol before generation of fluorescent cDNA, tissue removal, and imaging. We determined optimal tissue permeabilization time to be 16 min for both 2 and 7 DPI and therefore used this time for all tissues on the Spatial GEX slides. Tissue optimization slides were imaged following fluorescent RNA capture on the Axioscan at 20X under an HE DsRed filter cube. Images were exported as tiff files using Zeiss ZEN software for later use in spatial alignment. See **Supplementary Figure 1** for H&E images collected from all of the samples used in this study.

#### Visium FF Spatial GEX

Generation of spatial transcriptomic data was performed following the fresh frozen *Visium Spatial GEX User Guide* (Document number CG000239, Rev E, 10X Genomics). Tissue was permeabilized for 16 min after which RNA was released and mRNA is then bound by poly-adenylated tail capture to oligonucleotides on the capture spots (55 µm diameter) within the fiducial frames of the Visium GEX slide. After permeabilization, the mRNA was released and bound to oligonucleotides on the capture areas. This was immediately followed by reverse transcription, second-strand synthesis, denaturation, and cDNA amplification steps according to the manufacturer’s protocol using a T100 thermal cycler with the 10X Genomics slide adaptor. Reverse transcription was carried out on the thermal cycler for 45 min at 53°C and the slide was immediately removed to room temperature (RT) to perform second strand synthesis. 0.08 M KOH was freshly made just prior to the second strand synthesis from 8 M KOH (Acros Organics) as described in the user guide. The slide(s) were incubated in 0.08 KOH for 5 min at RT. Following incubation and another wash with Buffer EB (Qiagen), the second strand synthesis mix was added to the slide(s) and run through the second strand synthesis protocol on the thermal cycler for 15 min at 65 °C. After incubation, the slide(s) were washed with Buffer EB and then incubated in 0.08 KOH for 10 min at RT. During this time, 5 µL of Tris (1M, pH 7.0, Thermo Fisher) were added to the appropriate number of RNAse/DNAse-free PCR tubes on ice. After the 10 min incubation, each sample was added to its respective PCR tube and mixed 10 times by pipetting. Samples were then briefly centrifuged. Next, the PCR master mix was prepared as described in the user guide from the provided cDNA primer reagent, nuclease-free molecular water, and KAPA SYBR FAST qPCR mix. 9 µL of the mix was then added to the appropriate number of wells on a 96-well PCR plate plus an additional well for creation of a negative control. 1 µL of each sample was then added to their respective wells and mixed by pipetting 10 times. For a negative control, 1 µL of nuclease-free molecular water was instead added to the well.

The plate was then sealed and briefly centrifuged before performing PCR cycle number determination according to the user guide protocol to define the optimal number of cycles used for cDNA amplification (Cq). Cq estimation was performed with a C1000 Touch Thermal Cycler coupled to a CFX96 Touch Real-Time PCR Detection System (BioRad, Hercules, CA, USA): we found Cq to be 14-16 cycles for our experiments. cDNA amplification was then performed using the cDNA amplification mix (10X Genomics) on a T100 thermal cycler (BioRad) with appropriate cycle numbers set in the amplification protocol as described in the user guide. cDNA was then stored overnight at 4°C or at −20°C for ≈ 72 hours before performing cDNA cleanup. After quantitative PCR on the product, the Cq was determined, and the cDNA was amplified in order to have a sufficient input concentration into the library preparation process. The cDNA was also analyzed using the 12-channels Fragment Analyzer System from Agilent. The library preparation consisted of an initial fragmentation, end-repair and A-tailing step followed by size selected and cDNA amplification steps in order to incorporate the adaptors needed for sequencing.

cDNA cleanup/QC/quantification, library generation, and sequencing were performed by The American Genome Center at Uniformed Services University (Bethesda, MD, USA) following the user guide. Sample Index PCR cycles used for library construction were 10-14 cycles, which was determined based on our cDNA yield (14.5-139 ng) and % coverage of our tissue in the capture area (65-80%). Sequencing was performed on a NovaSeq 6000 platform (Illumina, San Diego, CA, USA) with 118 cycles as follows: 28 bp (Read 1), 10 bp (i7 Index), 10 bp (i5 Index), 90 bp (Read 2).

#### Space Ranger analysis pipeline

We deployed 10X Genomics Space Ranger 1.3.1 with resultant FASTQs and H&E tissue histology TIFF images to perform automatic slide image processing, read alignment, and spatial barcode/UMI counting. Reads were aligned to the Genome Reference Consortium Mouse Build 38 patch release 6 (GRCm38.p6). Space Ranger associates each read with a 55 μm spot within the histological tissue image based on the read’s spatial barcode to calculate per-spot gene expression.

#### Generation of GEX data from anatomical categories in Loupe Browser

Loupe Browser .cloupe files from each sample were then opened in 10X Genomics Loupe Browser v6.5.0. Loupe Browser was used to extract GEX data from spots in the tissue capture area of the left contused hemisphere from the following four anatomical locations based using the Allen Mouse Brain Atlas and Allen Reference Atlas – Mouse Brain (Allen Mouse Brain Atlas, mouse.brain-map.org and atlas.brain-map.org) as a reference and additional histological tissue landmarks from our H&E stained tissue images. (1) Cortex: to extract GEX data from the cortical area under the impact in CCI tissue and a respective control cortical area in Sham tissue, all grey matter spots above the white matter tracts of the Corpus Callosum (CC) and External Capsule (EC) were selected from the medial edge of the lateral ventricle to the lateral edge of the EC. (2) Corpus callosum-external capsule (CC-EC) white matter: All spots with at least 51% of their area within the CC or EC were selected. (3) anterior commissure (AC): All spots with at least 51% of their area within the AC were selected. (4) Striatum (STR): All spots with at least 51% of their area within the STR and housed below and medial to the CC-EC, above the AC, and not covering any part of the lateral ventricle or choroid plexus were selected.

#### Differential gene expression (DEG) analyses

Subsequent analyses were conducted in R (v4.2.0). We utilized the DESeq2 package (v1.36.0) following the recommended bulk RNAseq analysis workflow from the DESeq2 vignette. A nested experimental design in DESeq2 was used to test the group-specific temporal effects of injury and GR-1^+^ immune cell depletion on the brain tissue while controlling for individual sample variation. We also included a batch term per DESeq2’s recommendation to control for between batch variance. Genes where less than three samples had normalized counts less than or equal to 10 were filtered out, and the Wald test was used to test for significantly differentially expressed genes. We performed multiple-testing correction by the Benjamini-Hochberg procedure (target false discovery rate (FDR) = 0.05) and performed log_2_ fold change (LFC) effect size shrinkage as provided by the apeglm R package [29]. Genes with an absolute LFC ≥ + 1.0 or ≤ - 1.0 were considered biologically relevant, and DEGs with a *p*-adjusted value ≤ 0.05 were classified as significantly differentially regulated. Gene expression was compared between our eight experimental groups within a single anatomical category or in the case of our WM categories, the CC-EC and AC, both between groups in a single WM tract and also within a group between the CC and AC to identify DEGs.

#### Gene Set Enrichment Analysis (GSEA) Using Gene Ontology (GO) Pathway Terms

To identify enrichment for particular gene sets in our DEG results in response to injury and/or GR-1^+^ immune cell depletion, we performed GSEA via the R package clusterProfiler on selected experimental group and brain tissue comparisons [30, 31]. The Molecular Signatures Database (MsigDB) GO set (set number C5) and subset Biological Process was imported via the msigdbr R package (https://cran.r-project.org/package=msigdbr) for the species *Mus musculus* [32, 33]. Normalized enrichment scores were calculated and adjusted p values ≤ 0.05 were considered significantly enriched.

#### Performing Spatial Clustering and Analysis in Seurat

Since we identified that many genes in the CTX were expressed in a gradient away from the site of injury, we chose to perform spatial clustering analysis to identify differences in the amount of expression of injury-induced DEGs. We used the R package Seurat (v4.3.0.1) to normalize variance in gene counts across spots within a sample in our spatial GEX data and followed Seurat vignette guidelines for data analyses [34]. The transformed dataset then underwent integration and dimensionality reduction. We chose Harmony for integration of our dataset based on our experimental variables of Intervention (CCI/Sham surgery), Drug Treatment (Untreated, Control Ab, or anti-GR-1), Timepoint (2 or 7 DPI), and Batch [35]. The integrated dataset then underwent clustering with a resolution of 0.8 and the first 30 principal components were used for dimensionality reduction. Top markers of each cluster were identified along with changes to gene expression within clusters between experimental groups via DESeq2 bulk RNAseq analysis workflow as detailed above.

Spatial clustering analysis was performed on all tissue captured within the fiducial frame of the Visium slide. Occasionally, this included small medial portions of the contralateral cerebral hemisphere.

#### Experimental Design & Statistical Analysis

For confirmation of GR-1^+^ leukocyte depletion via flow cytometry, we employed a 4×2×2 study design such that CCI or Sham mice could have received either anti-GR-1 antibody or Control antibody until endpoints of either 2 or 7 DPI. For Visium Spatial GEX experiments, our study design resulted in eight experimental groups in which mice could have undergone either CCI or Sham procedures, received anti-GR-1, Control antibody, or no antibody treatment, and were sacrificed at either 2 or 7 DPI. For Visium GEX, each group had an n = 3-7 (see **Supplementary Materials, Table A**). DEGs in brain area pseudobulk analyses were identified by the Wald test with absolute LFC ≥ 1.0 considered biologically relevant and p-adjusted value ≤ 0.05 were classified as significantly differentially regulated. To identify DEGs in spatial clusters between groups, DEGs with p-adjusted values value ≤ 0.05 were classified as significantly differentially regulated. For GSEA, adjusted p values ≤ 0.05 were considered significantly enriched. Histogram data are presented as mean ± standard error of the mean (SEM).

#### Data Availability

Raw data files along with sample metadata is publicly available on Figshare at DOI: 10.6084/m9.figshare.27974352. Further sharing of data can be made upon request.

## Results

### Gene expression changes induced by CCI injury are principally segregated based on cytoarchitecture

Since TBI results from an initiating physical force whose impact acts on the brain tissue in a spatially-dependent manner, we aimed to characterize the transcriptional responses to injury and their spatial relationship to the initial insult in a well-characterized adult mouse model of mild CCI [24] at 2 and 7 DPI. Mice underwent CCI surgery or Sham procedures at 16-20 weeks-old, equivalent to 20-30-year-old humans [36, 37] and gene expression was mapped using the Visium Spatial GEX platform. Guided by the presence of the AC in coronal sections at the rostro-caudal focal point of impact, we compared transcriptional responses to injury in the grey matter territories of the CTX and STR with those of the white matter territories of the CC-EC and AC **[Fig. 1a and Supplementary Figure 2]**. Differential gene expression analyses were then performed by (pseudo)-bulk RNAseq between CCI and Sham groups across these structures at 2 and 7 DPI. DEG effect sizes suggest a topographical relationship between the breadth of gene expression changes and the structure’s distance from the site of impact, with total DEG number highest in the CTX and smallest in the AC at both 2 and 7 DPI **[Fig. 1b],** such that the deeper WM structure of the AC remained relatively unaffected by CCI. The exception to this is the STR at 2 DPI in which the total number of DEGs exceed those of the CC-EC closer to the site of impact, suggesting anatomical differences additionally influence the transcriptional response to injury. Plots of individual LFC values reveals larger and more widespread gene expression changes in upregulated compared to downregulated genes induced by CCI at both the acute 2 DPI and subacute 7 DPI endpoint **[Fig. 1c]**.

**Figure 1.**
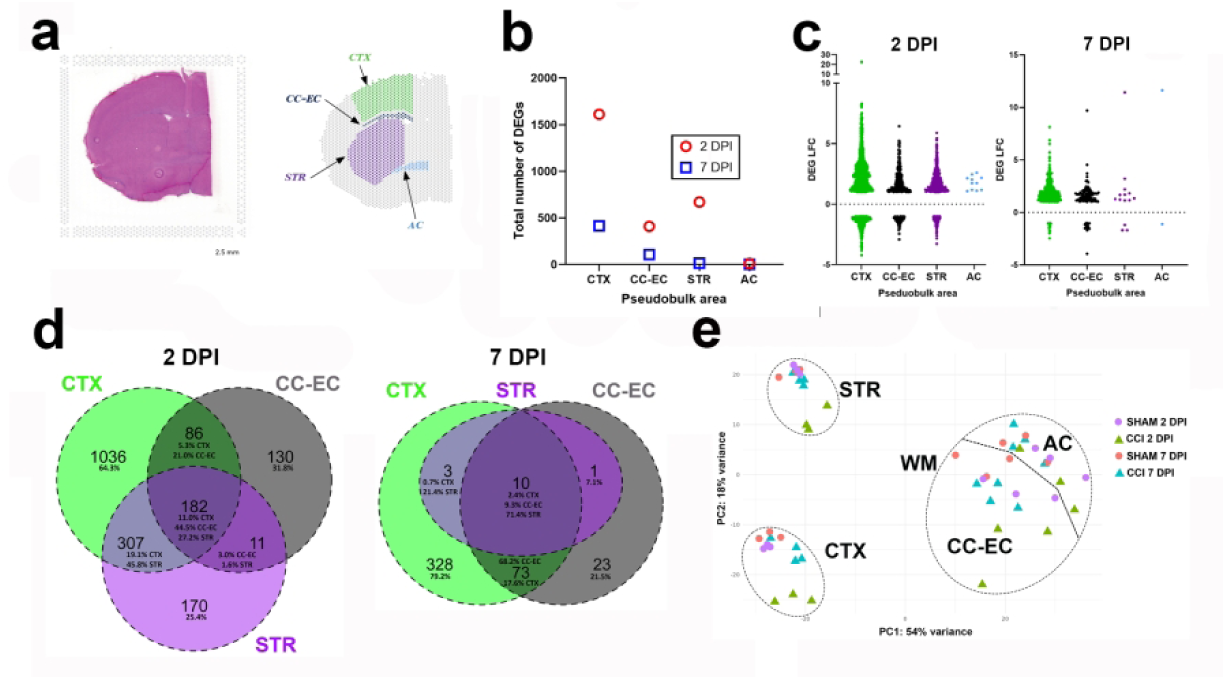
Spatial transcriptomics at the focal point of contusion following controlled cortical impact (CCI) in adult mice. **(a)** A representative brain hemi-section with hematoxylin and eosin (H&E) stain of the left hemisphere of the brain at the level of the AC (present at the level of bregma), the rostro-caudal point of impact at 2 mm left lateral from the midline. The image to the right is the brain hemi-section’s representative Loupe Browser spot selection for the 4 brain areas defined for (pseudo)-bulk RNAseq analysis, the cortex (CTX), corpus-callosum-external capsule (CC-EC), striatum (STR), and anterior commissure (AC). **(b)** Histogram of total number of CCI versus sham differential expressed genes (DEGs) identified in each anatomical territory at 2 and 7 days post-injury (DPI). **(c)** Scatter plots of the log2 fold-changes (LFC) of DEGs in each anatomical territory at 2 DPI and 7 DPI. **(d)** Venn diagram plots depicting the number of shared DEGs and the percentage (%) shared between the CTX, CC-EC, and STR at 2 DPI and at 7 DPI. **(e)** Principal component analysis (PCA) plot of all brain area samples shows clustering based on anatomical identity (PC1). Injury status results in further sub-clustering of the directly contused CTX samples along PC2. CCI is represented by ringed points, sham without this ring.

We next compared the number of unique DEGs per territory, DEGs that were shared across all three territories, and those differentially expressed within a single other territory. A total of 182 shared genes were differentially expressed in all 3 territories at 2 DPI and these accounted for 11% of all CTX DEGs, 45.8% of CC-EC DEGs, and 27.2% of STR DEGs **[Fig. 1d]**. These shared DEGs were highly enriched for genes with immune system function and involvement in the inflammatory response **[Supplementary Table 1]**. The CTX had the highest number (1036) and percentage (64.3%) of unique DEGs at 2 DPI and CTX DEGs were the highest percentage shared between either the CC-EC or STR alone (and the CC-EC and STR had the least DEG overlap). Although much reduced in magnitude, a similar pattern of DEG expression is observed amongst the 3 territories at 7 DPI with the highest number (328) and percentage (79.2%) of unique DEGs in the CTX, and CTX DEGs the highest percentage shared between either the CC-EC or STR alone **[Fig. 1d]**. Only 10 DEGs are shared across all three territories at this subacute endpoint and there are no DEGs unique to the STR **[Fig.1 d, Supplementary Table 1]**. Combined these data illustrate common but also topographically unique transcriptional responses to contusion injury in the three forebrain territories.

To better determine the relationship between the transcriptional response to injury with proximity to the site of impact and forebrain tissue anatomy, we next performed principal component analysis (PCA) on the pseudobulk DEG data. PCA showed that the anatomical/cellular identities of the different brain regions explained the majority of variance between samples, followed by injury status **[Fig. 1e]**. Comparing each sample’s gene expression in the white matter (WM of CC-EC or AC) and grey matter (GM of CTX or STR) demonstrates separation in PC1 (54% of variance) by WM or GM identity. PC2 (18% of variance) demonstrates separation of the GM territories based on proximity to the impact, such that the CTX samples, closer to the site of impact, clusters away from the more distal, ventral STR. Furthermore, there is clear separation of the 2 DPI CCI samples from the other CTX and STR samples in PC2, and weaker separation of the 7 DPI CCI CTX samples. In the WM, there is separation in PC2 of only the 2 DPI CC-EC samples. Taken together, PCA demonstrates the anatomical/cellular identities of the brain regions and secondarily the distance away from the focal impact and timepoint post-injury (for GM structures) drive gene expression changes within the contused forebrain.

### A cluster of injury-induced gene expression changes are shared across the cortex, striatum and corpus-callosum-external capsule

We next analyzed the 182 DEGs shared across all three territories at 2 DPI. Remarkably, the directionality of shared DEG gene expression changes was the same in each territory with 174 DEGs upregulated by CCI and only eight genes downregulated (*AI593442*, *Bcan*, *Cldn10*, *Cyp4f15*, *Jph1*, *Ppp1r3g*, *Prkcg*, *Ptgds*) **[Supplementary Table 1]**. Genes highly upregulated in all three territories at 2 DPI include neuroinflammatory and wound healing genes such as the protease inhibitor and cytokine transporter *A2m*, the chemokine *Ccl2*, the matrix metalloproteinase (MMP) inhibitor *Timp1*, the serine protease inhibitor *Serpina3n*, the secreted phosphoprotein *Spp1*, the calcium-dependent enzyme transglutaminase 1 (*Tag1*), and the microtubule protein *Tubb6*.

### Interferon signaling and leukocyte-mediated immunity gene pathways are acutely upregulated in the contused cortex

The largest transcriptional response to injury at 2 DPI is in the CTX with 1611 total DEGs, 1128 of which are upregulated. Significantly upregulated genes in the CTX include multiple chemokines (e.g. *Cxcl10, Cxcl16, Ccl2, Ccl3, Ccl6, Ccl7, Ccl12*), the cytokine *IL33*, complement genes (*C1qa, C1qb, C1qc, C4b*), and numerous interferon-stimulated genes (e.g. ISGs *Fcgr1*, *Fcgr4*, *Fcgr2b*, *Ifi207*, *Ifi204*, *Irf7*, *Irf8*, *Irf9*), and other markers of immune and inflammatory responses such as *Cd9, Cd44, Lgals3,* and *Tnfrsf12a* **[Fig. 2a, Supplementary Table 1]**. Of these, upregulation of *Ccl3*, *Ccl6*, *Ccl7* and *IL33* is unique to the CTX amongst the three territories at 2 DPI. Consistent with the large representation of upregulated interferon-response genes in the CTX, the most positively enriched Gene Ontology (GO) Biological Process pathways identified via Gene Set Enrichment Analysis (GSEA) include *response to interferon gamma* and *response to type I interferon* **[Fig. 2b, Supplementary Table 2]**. GO terms related to leukocyte cell chemotaxis, as well as to leukocyte cell activation and cellular immunity are highly positively enriched in the 2 DPI CTX as well **[Fig. 2b, Supplementary Table 2]**. Overall, GSEA confirms innate immunity gene pathways are robustly upregulated in the CTX at 2 DPI.

**Figure 2.**
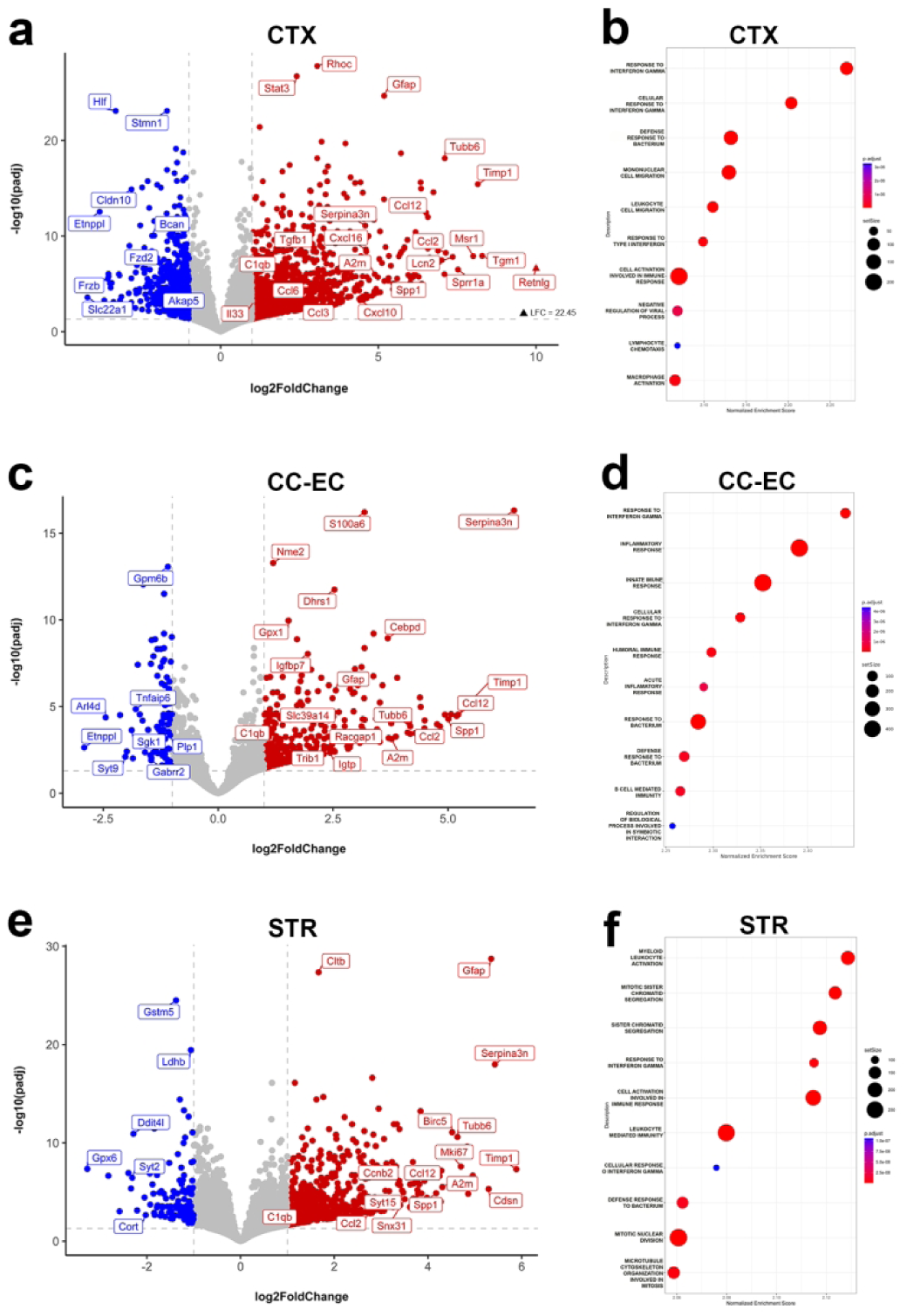
Interferon signaling and leukocyte-mediated innate immunity gene pathways are acutely upregulated by CCI. **(a)** Volcano plot of significantly (p-adj < 0.05) upregulated (LFC > +1.0, red) and downregulated (LFC < −1.0, blue) DEGs in the CTX at 2 DPI. **(b)** Dotplot of the 10 most positively significantly enriched Gene Ontology (GO) Biological Process (BP) terms in the CTX at 2 DPI by normalized enrichment score (NES) identified via gene set enrichment analysis (GSEA). The size of the dot indicates the set size of the enriched GO term and the color of the dot indicates the relative p-adj value (blue = high/less significant, red = low/more significant). **(c)** Volcano plot of significant DEGs and **(d)** dotplot of enriched GO terms by GSEA in the CC-EC at 2 DPI. **(e)** Volcano plot of significant DEGs and **(f)** dotplot of enriched GO terms by GSEA in the STR at 2 DPI.

Genes downregulated by CCI at 2 DPI were largely unique to each structure. Of the 483 downregulated DEGs in the CTX at 2 DPI, only 83 (17%) were shared with either the CC-EC, STR or both. Downregulated genes unique to the CTX at 2 DPI included genes functioning in synaptic transmission (*Adcy8*, *Camkv*, *Gria3*, *Grin2c*, *Slc6a1)* and in neuronal differentiation (*Akap5*, *Frzb*, *Fzd2*, *Stmn1*) **[Fig. 2a]**. This is reflected in the GSEA of the most negatively enriched GO terms that include *synapse assembly*, *ligand-gated ion channel signaling pathway*, and *dendrite morphogenesis* **[Supplementary Table 2]**.

### Shared immune and inflammatory gene pathways are acutely activated in the injured neocortex and corpus-callosum-external capsule

Gene expression changes in the CC-EC largely overlap those of the CTX following CCI at 2 DPI: the CC-EC shares 66% of its DEGs with the CTX. There is a strong representation of genes involved in neuroinflammatory and immune processes following injury in the CC-EC, for example the Ccr2 ligands *Ccl2* and *Ccl1* are robustly upregulated, as are *Cd9*, *Cd44*, *Irf7*, *Irf9*, *Serpina3n* and *Spp1* **[Fig. 2c]**. Uniquely upregulated genes in the CC-EC included the ISG GTPase *Igtp*, the divalent metal ion transporter *Slc39a14*, the serine/threonine pseudokinase *Trib1*, and the GTPase-activating protein *Racgap1* **[Fig. 2c, Supplementary Table 1]**. Similar to the CTX, the highest positive GO term identified by GSEA in the CC-EC is *response to interferon gamma* and highly positive GO terms are related to innate immune and inflammatory responses **[Fig. 2d, Supplementary Table 2]**. Specific to the CC-EC was the relatively strong positive enrichment by GSEA of gene sets involved in the processing and presentation of antigens at 2 DPI [see **Supplementary Table 2]**. In general, the large number of shared DEGs between the CTX and CC-EC is reflected in shared activated gene set pathways with 14 of the highest enriched GO terms shared between the territories. Downregulated DEGs unique to the CC-EC at 2 DPI include the neutrophil regulator *Sgk1,* the membrane-associated GTP-binding protein *Arl4d* (shown to regulate neurite outgrowth [38]), the GABA receptor subunit *Gabrr2*, and the myelin protein *Plp1*. As was the case in the CTX, negatively enriched GO terms such as *ligand-gated ion channel signaling pathways* and *glutamate receptor signaling pathway* are consistent with a suppression of neuronal transmission in the injured CC-EC at 2 DPI.

### Gene pathways involved in the cell cycle and mitosis are acutely activated in the injured striatum

The number of injury-induced DEGs in the STR is larger than that in the CC-EC at 2 DPI, despite this structure being more distal to the contusion. STR gene expression changes are similar to the CTX in that they are enriched for genes involved in innate immune and inflammatory responses such as the chemokines *Ccl2*, *Ccl8*, *Cc112*, and interferon response genes. Highly upregulated STR DEGs shared with both the CTX and CC-EC include *Serpina3n*, the inhibitor of apoptosis (IAP) gene *Birc5*, and the myeloid cell pro-inflammatory, pro-survival gene *Bcl3* **[Fig. 2e]**. Overall, however, there is greater overlap of STR DEGs with those of the CTX than those of the CC-EC. As expected, gene sets involved in leukocyte mediated immunity and ISGs are highly positively enriched in the STR at 2 DPI **[Fig. 2f]**. A distinct injury feature of the STR at 2 DPI is the strong positive enrichment for gene pathways for chromatid segregation and mitotic cell division **[Fig. 2f, Supplementary Table 2]**. Mitosis cell cycle genes robustly upregulated in the STR include *Mki67*, *Birc5*, *Bcl3*, and *Ccnb2* **[Fig. 2e]**. DEGs uniquely downregulated in the STR at 2 DPI include the chemokine scavenger receptor *Ackr2*, *Ddit4l*, a negative regulator of mTOR and innate immune activity [39], the synaptotagmin protein *Syt2*, and the neuropeptide cortistatin, *Cort* **[Fig. 2e]**. GSEA indicates suppressed neural activity in the STR at 2 DPI with GO terms *ligand-gated ion channel signaling pathways* and *glutamate receptor signaling pathway* negatively enriched at 2 DPI.

### Interferon gamma signaling and adaptive immunity gene pathways are activated subacutely

At 7 DPI there is a significant reduction in injury-induced gene expression changes across all three brain territories **[Supplementary Table 1]**. Only ten DEGs are shared across all three territories at 7 DPI and, as was the case at 2 DPI, the directionality of shared DEG gene expression changes were the same in each territory with 9 DEGs upregulated and only the non-coding RNA *Gm47283* downregulated in the CTX, CC-EC and STR. The nine shared upregulated DEGs include some of the highest injury-induced increases in gene expression at 7 DPI measured by LFC such as *A2m*, *C4b* complement, and the reactive astrocyte markers *Gfap* and *Lgals3bp* **[Fig. 3a, c]**.

**Figure 3.**
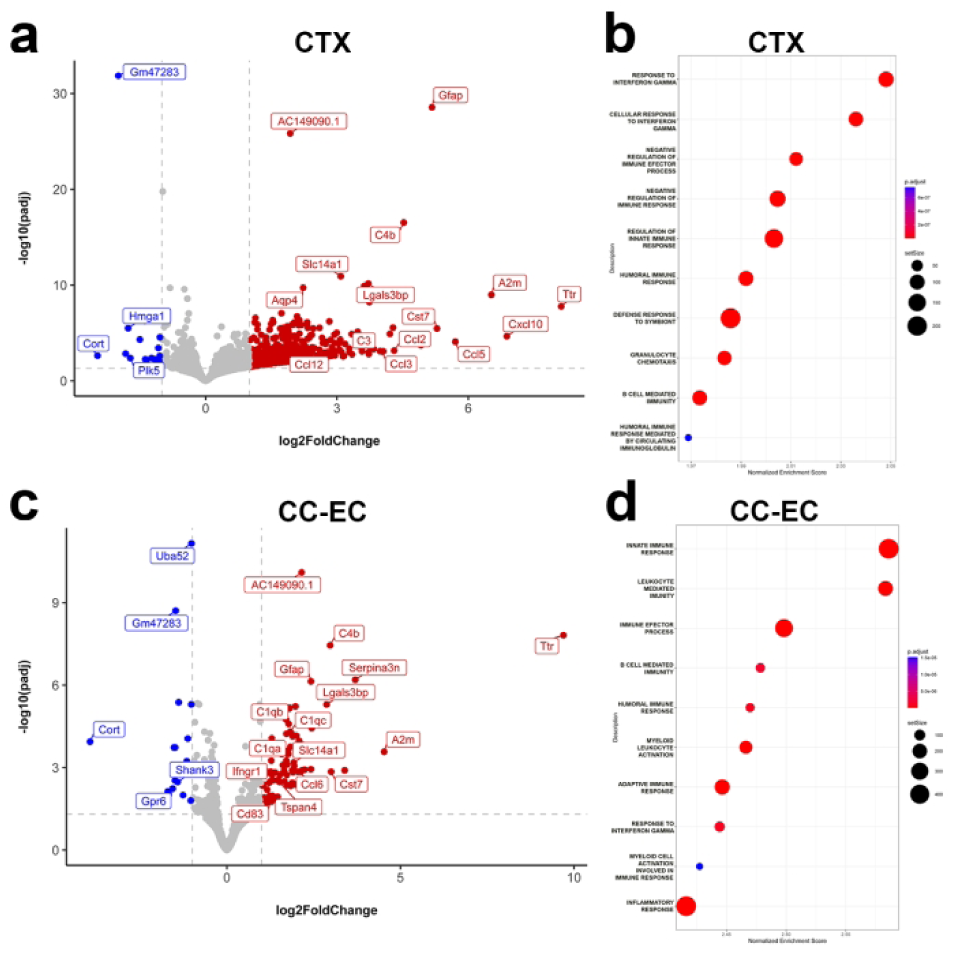
Adaptive immunity gene pathways are activated subacutely in CCI mice. **(a)** Volcano plot of significantly (p-adj < 0.05) upregulated (LFC > +1.0, red) and downregulated (LFC < - 1.0, blue) DEGs in the CTX at 7 DPI. **(b)** Dotplot of the 10 most positively enriched Gene Ontology (GO) terms in the CTX at 7 DPI by normalized enrichment score (NES) identified via gene set enrichment analysis (GSEA). The size of the dot indicates the set size of the GO term and the color of the dot indicates the relative padj value (blue = high, red = low). **(c)** Volcano plot of significant DEGs and **(d)** dotplot of enriched GO terms by GSEA in the CC-EC at 7 DPI.

There are 414 injury-induced DEGs by 7 DPI in the CTX, down by ∼ 75% from the number at 2 DPI, and the vast majority of the DEGs at 7 DPI are upregulated in the CTX (395 of the 414). Strongly upregulated genes in the 7 DPI CTX include the bone marrow stromal cell antigen *Bst2*, chemokines *Ccl2*, *Ccl3*, *Ccl5*, *Ccl12*, *Cxcl10*, complement *C3*, and the cysteine protease inhibitor *Cst,* consistent with continued innate immune system activity **[Fig. 3a]**. Of these, *Ccl5*, *Cxcl10* and *C3* are uniquely upregulated in the CTX at 7 DPI suggesting region-specific functions. Interferon-related genes remain strongly upregulated at 7 DPI and include the type I interferon genes *Irf7* and *Isg15* [40] and the microglial inflammation modulator *Ifi27l2a* **[Supplementary Table 1]**. The sustained upregulation of interferon signaling genes is reflected by GSEA that again identifies *response to interferon gamma* as the most positively enriched GO term in the injured CTX **[Fig. 3b]**. Other strongly positively enriched GO terms reveal the transcriptional activation of adaptive immunity gene sets in the CTX by one-week post-injury and include *humoral immune response* and *B cell mediated-immunity* **[Fig. 3b]**. Robustly upregulated genes with humoral B-cell functions include *A2m*, the B-cell regulator *Bst2*, the MHC class I antigen *H2-Q7*, and the lymphocyte cell surface receptor *Cd84* **[Fig. 3a, Supplementary Table 1]**. A temporal shift in the immune response in the cortex over the first post-injury week is further supported by the positive enrichment of GO terms for the negative regulation of the immune response in the 7 DPI CTX **[Fig. 3b]**. Genes specifically downregulated in the CTX at 7 DPI include *Cort* and the kinase-regulator *Plk5* linked to synaptic homeostasis [41] **[Fig. 3a]**.

In the CC-EC by 7 DPI, there are 117 injury-induced DEGs following CCI. As was the case at 2 DPI, many of the innate inflammatory genes upregulated in the CTX are similarly upregulated in the CC-EC at 7 DPI. These include *Bst2*, *Ccl3*, *Cst7*, *Ccl9*, and *C1q* complement genes **[Fig. 3c]**. The chemokine *Ccl6* is one of the limited numbers of injury-induced genes unique to the CC-EC **[Fig. 3c]**, suggesting a distinct role for this Ccr1 ligand in the WM response to injury at 7 DPI. Other upregulated DEGs unique to the CC-EC include the interferon gamma receptor *Ifngr1*, the immunomodulatory protein *Cd68*, and the cell surface glycoprotein *Tspan4*. GSEA of the 7 DPI CC-EC reveals largely the same pathways activated in the 7 DPI CTX with continued activation of *innate immune response* genes (the highest positively enriched GO term) plus GO terms *B cell mediated-immunity*, *humoral immune response* amongst others indicative of the activation of adaptive immunity **[Fig. 3d]**. Downregulated DEGs specific to the CC-EC at 7 DPI include the orphan G-protein coupled receptor *Gpr6* and the synaptic scaffold *Shank3*. Shared among the few downregulated DEGs in the CTX (19 DEGs) and CC-EC (15 DEGs) at 7 DPI are *Cort* and *Camk2n1*, a calcium/calmodulin-dependent protein kinase II inhibitor **[Fig. 3c, Supplementary Table 1]**. The STR has a very small transcriptional response to injury at 7 DPI with only 14 DEGs and no positively enriched pathways by GSEA. The highest upregulated genes in the STR by LFC are the nine genes upregulated in all three territories at 7 DPI **[Supplementary Table 1]**. Taken together the 7 DPI data indicate a transition from innate to adaptive immune signaling in the dorsal CTX and CC-EC, and a significant resolution of injury-induced gene expression changes in the more ventral STR within the first week post-injury.

### Spatial clustering identifies an evolving injury-induced gene expression cluster in the cortex of CCI mice at 2 DPI and 7 DPI

To allow for an unbiased examination of the spatial transcriptional responses to CCI, and because brain tissue areas have cellular sub-regions not captured by our pseudobulk comparisons, we next performed spatial clustering analysis across all of our tissue samples. Visualization of our spatial cluster results by uniform manifold approximation and projection (UMAP) demonstrates the gene expression data clusters into 29 groups that can be mapped onto representative tissue slices **[Fig. 4a, Supplementary Figure 3]**. Spatial clustering reveals two injury clusters only present in CCI mice, clusters 6 and 13. Cluster 6 is the injury cluster at 2 DPI encompassing the outer layers of the primary somatosensory cortex, portions of the more medial primary motor and cingulate cortex, and deeper cortical layers at the focal point of impact. **[Fig. 4a]**. Indeed, cluster 6 entirely replaces cluster 10 found in the outer cortical layers of sham mice [see **Fig. 4a]**. **Supplementary Table 3** lists cortical cluster genes, including cluster 6, and their fold-change increase versus all other 2 DPI CCI spatial clusters. Cluster 6 marker genes are related to the inflammatory response, cytokine and chemokine production, and receptor binding. They include the related interleukins *Il6, Il9*, and *Il11*, chemokines *Cxcl1*, *Ccl2*, *Ccl7* and receptors *Ccr2* and *Ccr7*. GSEA of cluster 6 markers reveals the enrichment of antimicrobial humoral response peptides (the innate response to alarmins), mononuclear and lymphocyte migration, and inflammatory response gene pathways in this CCI-induced cluster **[Fig. 4b]**.

**Figure 4.**
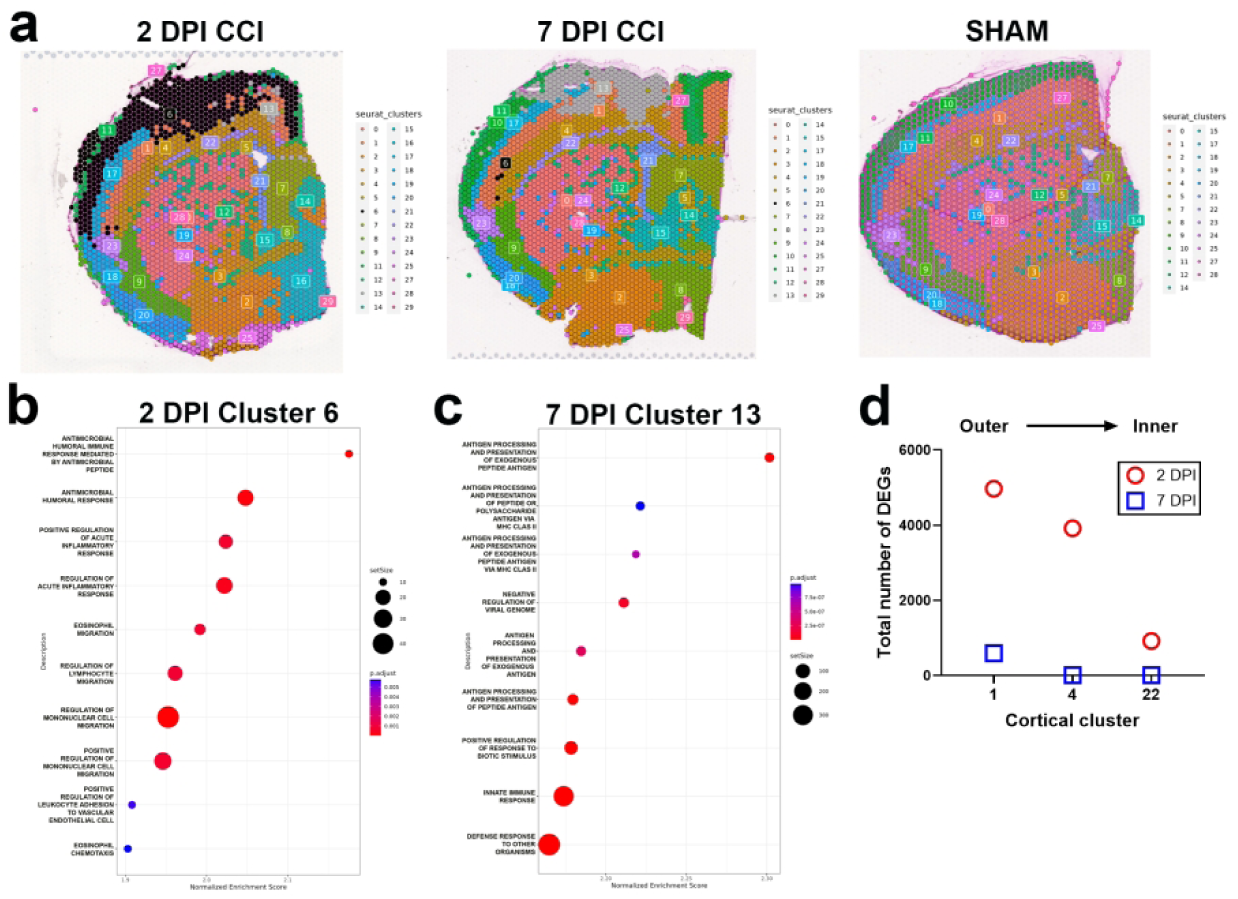
Spatial clustering identifies injury-induced gene expression clusters in the neocortex of CCI mice at 2 DPI and 7 DPI. **(a)** Representative forebrain tissue slices with UMAP projections of spatial cluster identities. Dotplot of enriched GO terms by GSEA in **(b)** the injury cluster 6 at 2 DPI and **(c)** the injury cluster 13 at 7 DPI. The size of the dot indicates the set size of the GO term and the color of the dot indicates the relative padj value (blue = high, red = low). **(d)** Histogram of the number of DEGs per cluster in the cortex of mice at 2 DPI and 7 DPI.

At 7 DPI, cluster 6 is replaced by cluster 13 as the injury cluster. When compared to cluster 6 at 2 DPI, cluster 13 is smaller in size, both in medio-lateral and dorso-ventral extent, and no longer encompasses significant portions of the deeper layers of the primary somatosensory cortex **[Fig. 4a]**. Spatial contraction of the injury cluster in the contused cortex between 2 and 7 DPI is matched by the renewed presence of cluster 10 within the outer cortical layers [see 7 DPI CCI tissue cluster map in **Fig. 4a]**. These changes in cortical clustering signify active tissue recovery between 2 and 7 DPI. As was the case in our pseudobulk analysis, genes uniquely enriched in cluster 13 support the activation of adaptive immunity and antigen presentation gene pathways in addition to innate immune responses in the injured tissue at 7 DPI **[Supplementary Table 3]**. Gene markers of cluster 13 include *Cd69,* which is expressed in activated T cells, the T cell modulatory chemokine *Cxcl9* [42], and the B cell lymphocyte chemoattractant chemokine *Cxcl13*. Accordingly, gene pathways for antigen processing and presentation are the most positively enriched in cluster 13 by GSEA **[Fig. 4c]**. Also present in cluster 13 are multiple genes associated with homeostatic and disease-associated microglia (DAM) cell states, for example the homeostatic genes *Aif1* (Iba1), *Fcrls*, *P2ry6*, and DAM markers *C5ar1*, *Irgm1* and *Fcgr1* **[Supplementary Table 3]**.

Although the spatial resolution of clustering did not recapitulate anatomically the six histologically defined layers of the CTX, clustering did segregate the CTX into layers stacked across the axis of the outer to inner CTX. Thus, cortical clusters present in both CCI and Sham mice were spatially distinguished by their proximity to the point of impact. We took advantage of this feature to examine the relationship between cortical depth and the transcriptional response to injury by comparing CCI versus sham DEGs between layers. We focused on cortical clusters 1, 4 and 22. These clusters were aligned superiorly to inferiorly across the full medio-lateral extent of impact, and were present in both CCI and sham mice at the 2 and 7 DPI endpoints (although also aligned superiorly to inferiorly, cluster 17 present in both CCI and sham mice does not extend into the medial impact zone, and cluster 10 is not present in CCI mice at 2 DPI). At 2 DPI we found a positive relationship between the number of CCI versus sham DEGs within these clusters and the proximity of the cluster to impact: DEG number was highest in the outer-most cluster, cluster 1, and lowest in the inner-most cluster 22 **[Fig. 4d]**. These results reveal a direct relationship between acute injury-induced gene expression changes and proximity to impact within the laminar structured CTX.

### Modified transcription of genes with synaptic, metabolic and mRNA transcription-translation functions with GR-1^+^ immune cell depletion

As neutrophil and monocyte infiltration into the injured mouse brain peaks during the first week post-injury, we sought to determine the spatial effects of these infiltrating cells on injury-induced gene expression changes in the brain with the hypothesis that regions more proximal to the site of impact would have the strongest predisposition to signaling by infiltrating immune cells. We used an antibody-mediated approach to deplete circulating leukocytes in CCI and sham groups, beginning the day before surgery and continuing until the 2 or 7 DPI endpoint. Mice were administered monoclonal IgG2b rat anti-mouse GR-1 antibody or an isotype rat monoclonal IgG2b control antibody that is non-reactive in mice as described previously in a mouse model of intracerebral hemorrhage [43]. Blood and spleen tissues were analyzed by flow cytometry at 2 and 7 DPI. Anti-GR-1 treatment resulted in a more robust depletion of Ly6G^+^ and a modest depletion of Ly6C^+^ immune cells at both timepoints in the periphery **[Supplementary Figure 4],** which is consistent with this particular clone’s (RB6-8C5) epitope reactivity [25, 44, 45]. Weight measurements made throughout the timeline of treatment demonstrated that anti-GR-1 treatment resulted in a small but significant reduction in weight in CCI and sham mice when compared to those treated with control antibody after 6 days of treatment **[Supplementary Figure 5]**.

Next, we determined if the depletion of circulating GR-1^+^ immune cells resulted in the attenuation of infiltrating GR-1^+^ immune cells into the ipsilateral contused or sham (left) brain hemisphere at 2 and 7 DPI following anti-GR-1 treatment. Within our flow cytometry panel, we included markers of other immune cell populations (see **Supplementary Materials, Table B**) to segregate central microglia from infiltrating immune cells through CD45 expression levels and inclusion/exclusion makers for myeloid cell subsets [see **Supplementary Figure 6** for gating strategy]. Flow cytometry confirmed anti-GR-1 treatment reduced the number of infiltrating Ly6G^+^ neutrophils and Ly6C^low-int^ monocytes at both timepoints; however, infiltrating Ly6C^hi^ monocytes were unaffected by anti-GR-1^+^ treatment at 2 DPI but experienced reduced infiltration at the later endpoint of 7 DPI **[Supplementary Figure 7].**

To identify the molecular effects of reduced GR-1^+^ immune cell infiltration in the injured brain, we used our pseudobulk approach to compare gene expression changes in CTX, CC-EC and STR territories between anti-GR-1 or control antibody treated CCI mice at 2 and 7 DPI. Anti-GR-1 treatment resulted in minor gene expression changes in the CTX, CC-EC and STR with DEGs shared across territories **[Supplementary Table 4]**. *AC149090.1*, an ortholog of human *PISD*, a mitochondrial phospholipid decarboxylase linked to autophagy [46, 47], is upregulated in the CTX, CC-EC and STR at 2 DPI, and *Lrrc1*, a gene of unknown function associated with cancer progression and tumor immune cell infiltration [48], is downregulated in all three territories at 7 DPI. (Ttr downregulation detected in all three territories at 2 DPI, see **Supplementary Table 4,** is likely an artefact due to sample differences in the inclusion of the choroid plexus, the sole site of Ttr expression in the brain [49]). Consistent with limited transcriptional effects of GR-1^+^ immune cell depletion, PCA separated the pseudobulk DEG data by anatomical region but not by antibody treatment **[Supplementary Figure 8]**. GSEA of the pseudobulk data revealed the positive enrichment of multiple GO terms for central nervous system development, primarily synapse and dendrite morphogenesis and function, in the CTX and STR of GR-1^+^ immune cell-depleted mice at 2 DPI **[Figure 5a, b, Supplementary Table 5].** There were no enriched GO terms in the CC-EC at 2 DPI or in any of the pseudobulk territories at 7 DPI.

**Figure 5.**
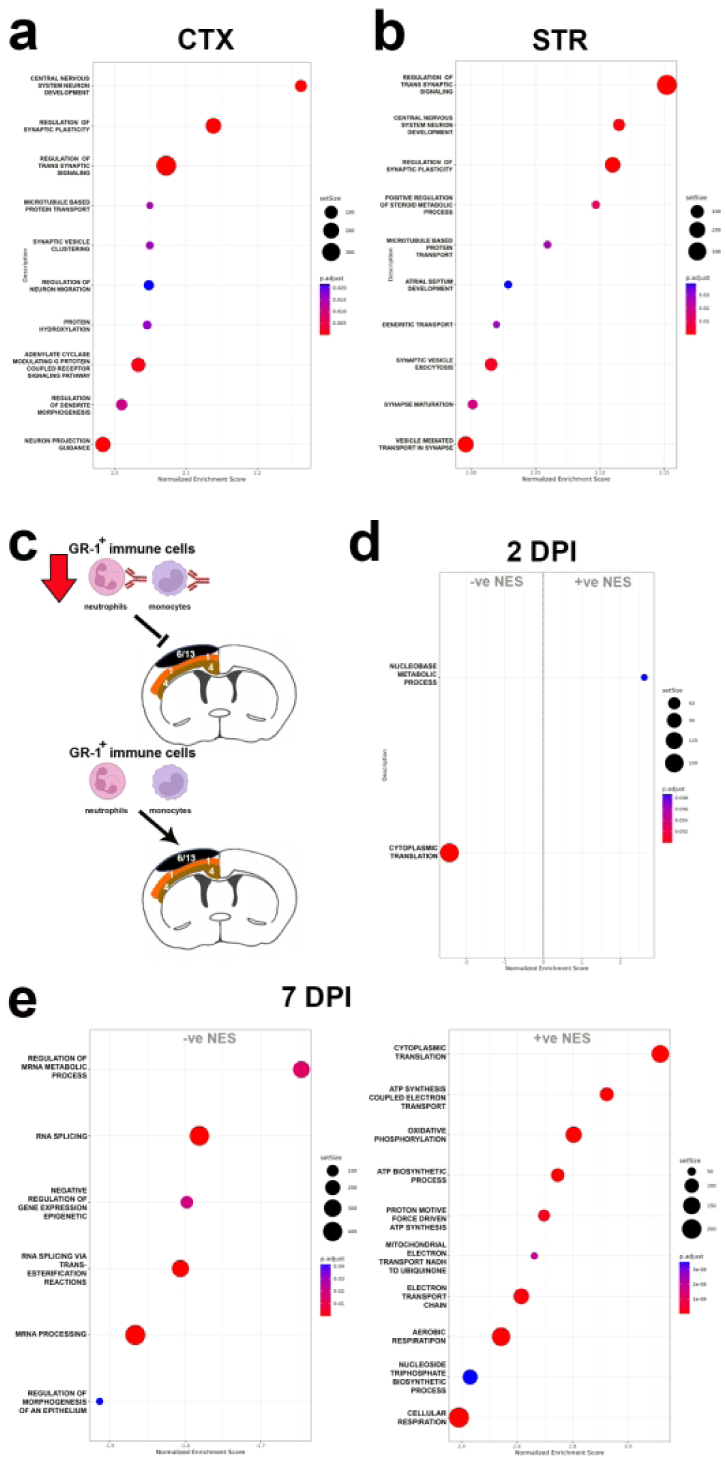
GR-1^+^ immune cell depletion results in acute gene expression changes in the injured neocortex and striatum. Dotplot of positively enriched GO terms by GSEA in **(a)** the CTX and **(b)** the STR at 2 DPI. **(c)** Schematic of GR-1+ immune cell depletion and spatial clusters located in mid and deeper layers of the injured neocortex. **(d)** Dotplot of negatively enriched GO terms reveals downregulation of *cytoplasmic translation* in cluster 1 in the neocortex at 2 DPI. **(e)** Dotplots of negatively (left) and positively (right) enriched GO terms in cluster 1 at 7 DPI reveals the downregulation of mRNA processing and metabolism gene pathways, and the upregulation of *cytoplasmic translation* and gene pathways functioning in ATP synthesis, the electron transport chain, and oxidative phosphorylation. The size of the dot indicates the set size of the GO term and the color of the dot indicates the relative padj value (blue = high, red = low).

Since unbiased spatial clustering revealed a relationship between differential gene expression and proximity to the physical impact in the neocortex of CCI mice, we repeated this analysis on anti-GR-1 and control IgG treated samples. Visualization of this data by UMAP similarly segregated the data into 29 groups including injury clusters 6 and 13 present at 2 and & DPI respectively **[Supplementary Figure 9]**. However, we found no significant changes with GR-1^+^ immune cell depletion in injury clusters 6 or 13 of the CTX: instead, GR-1^+^ immune cell depletion effects were detected in the mid-cortical territory of cluster 1 at both endpoints **[see Figure 5c]**. Cluster 1 gene expression changes were small with LFCs < ± 1 but GSEA revealed GO terms significantly altered with GR-1^+^ immune cell depletion. These included the negative enrichment for the GO term *cytoplasmic translation* at 2 DPI and bi-directional effects at the later 7 DPI endpoint **[Fig. 5d, e and Supplementary Table 5]**. At 7 DPI, GO terms related to mRNA metabolism, splicing and processing were negatively enriched, whereas *cytoplasmic translation* and GO terms related to ATP synthesis, the electron transport chain, and oxidative phosphorylation were positively enriched **[Fig. 5e]**. Interestingly, we did not detect anti-GR-1 treatment effects in cluster 4, located immediately deeper to cluster 1 in the cortex **[see Fig. 5c]**, indicating a discrete spatial response to GR-1^+^ immune cell depletion in the contused cortex.

In conclusion, transcriptional responses to injury in the first post-injury week in CCI mice are dominated by interferon gamma (IFN-γ) signaling and leukocyte-mediated immunity across the CTX, CC-EC and STR. However, there exists key spatiotemporal differences. These include the finding that anatomical/cellular identities of different brain regions are the primary driver of divergent transcriptional effects, but also the observation that within regions of relatively similar cytoarchitecture, such as within the cortex, there is a positive relationship between proximity to the point of impact and the size of the transcriptional response. A temporal feature, consistent across all three anatomical territories, is the transition from acute granulocyte-mediated, innate immune transcriptional responses toward the activation of gene sets functioning in B cell-mediated, humoral immunity by the end of the first post-injury week. Lastly, we identify a novel transcriptional feature of depleted GR-1^+^ immune cell infiltration that suggests neutrophil and monocyte signals contribute to multifaceted transcriptional responses in injured brain tissue to influence synaptic, metabolic, and mRNA transcription and translation in the brain post-injury.

## Discussion

In this study we hypothesized that the transcriptional landscape in the contused brain tissue would have a spatial relationship with the initial physical insult. Additionally, since clinical sequelae appear dependent upon the involvement of discrete brain structures such as the WM tracts [50–54], we further expected that contusion would produce differential gene changes in the WM versus GM structures in the brain. We first employed a (pseudo)bulk RNAseq analyses in which the major anatomical areas underneath the site of impact, the CTX, CC-EC, STR, and AC were delineated via histological landmarks. Comparisons of gene expression within these structures between CCI and sham mice revealed significant DEGs in the CTX, STR, and CC-EC but not the AC. The directly contused CTX had the highest number of DEGs, as expected based on clinical and preclinical research that the severity of symptoms is correlated to the severity of the brain injury [55]. However, what was surprising to see in our CCI model is that tissue topography/cytoarchitecture drove more of the differences between brain structures than did injury, likely due to the relatively mild nature of our low-speed and low-depth contusion CCI model. Interestingly, although individual DEGs were spatially divergent, gene pathways activated by CCI were largely shared across the CTX, STR, and CC-EC.

### IFN-γ signaling responses in the contused forebrain

A consistent feature of our data was the widespread and prominent transcriptional upregulation of ISGs in response to CCI injury and the common activation of IFN-γ signaling responses in the CTX, CC-EC and STR. While IFN-γ is classically associated with adaptive immunity, it also plays significant roles in acute immune responses, particularly in CNS injury and inflammation [56, 57]. IFN-γ is secreted by monocytes, T cells and microglia [58, 59], and IFN-γ signaling regulates leukocyte trafficking at brain borders (including at the choroid plexus) to control leukocyte infiltration [60–62]. Chemokines and cytokines (C/Cs) acutely upregulated by CCI in the forebrain such as *Cxcl10, Ccl12, Ccl2,* and *Ccl3* are activated by IFN-γ, along with IFN-sensitive factor genes such as *Ifitm3*, *Irf8* and *Irf9* detected in our mice. Later, *Cxcl9*, an IFN-γ induced chemoattractant for T lymphocytes and Natural Killer (NK) cells [63], is prominently upregulated in the injury cluster (cluster 13) of the CTX at 7 DPI.

In the context of TBI, IFN-γ signaling is thought to be important for microglia priming, the activation of astrocytes, and recruitment of T cells during the first week post-injury [64, 65]. In particular, IFN-γ signaling likely skews CD4^+^ T helper cells towards a pro-inflammatory subset, Th1, implicated in poor cognition following injury [19, 66, 67], likely due to its anti-proliferative signaling properties extending beyond immune cells to neurons [68]. Despite the common detection of IFN-γ signaling by GSEA in all three regions at 2 DPI, our data supports divergent downstream signaling effects in the CTX, CC-EC and STR. For example, the transcription factor *Irf8* capable of amplifying type II and type I IFN responses in immune cells [69] is upregulated in the CTX and STR but not CC-EC. In contrast, the type II interferon immunity-related GTPase *Igtp* was uniquely upregulated in the CC-EC white matter at 2 DPI in our CCI mice. Igtp has well described functions in antimicrobial resistance and the clearance of pathogens [70] but may exert anti-apoptotic effects in neurons following intracerebral hemorrhage [71].

### Chemokine and cytokine signaling

Other neuroinflammatory genes acutely upregulated in our study include proinflammatory chemokine and cytokines (C/Cs) previously linked to TBI such as *Ccl2*, *Ccl3*, *Ccl5*, *Ccl7*, *Ccl12* and *Cxcl10* [24, 72]. While the role of Ccl2 in the recruitment of inflammatory Ccr2^+^ myeloid cells to the injured brain is established [16, 72–75], Ccl7 and Ccl12 can exert many of the same functions as Ccl2 by signaling through the same Ccr2 receptor [72]. Indeed, *Ccl2* and *Ccl12* are the two most upregulated chemokines in the CTX and CC-EC with CCI at 2 DPI. The redundancy of signaling through Ccr2 following CCI may be important for the pathophysiology of secondary injury, particularly to recruit pro-inflammatory Ccr2^+^ myeloid cells [15, 76–78]. For example, *Ccl12* deletion does not have a detectable impact on monocyte trafficking under homeostatic conditions [79] and yet upregulation of *Ccl12* has been identified as a possible driver of increased Ccr2^+^ monocyte infiltration and exacerbated memory dysfunction in aged mice subjected to a TBI [80]. Similarly, cell-specific differences in the expression of Ccr2 ligands likely contribute to regionalized differences in pathology. For example the Ccr2 ligand *Ccl8* [81], shown to drive infiltration of cytotoxic CD8^+^ T cells in radiation-induced and ischemia-induced brain injury along with Ccl2 [82], is robustly upregulated with *Ccl12* and *Ccl2* in the STR at 2 DPI.

In addition to C/Cs well-validated in the field, we detected upregulation of the cytokine *Il33* in the contused CTX at 2 DPI consistent with its role as an alarmin with broad pro- and anti-inflammatory effects in astrocytes, oligodendrocytes, microglia and T helper cells in the CNS [83]. Also, acutely upregulated in the CTX, STR, and to a lesser extent the CC-EC, was the chemokine *Cxcl16*. The role of *Cxcl16* in TBI pathology is unknown but the encoded protein is reported to be downregulated compared to shams in a proteomic study of mild experimental TBI in mice [84]: Cxcl16 protein levels were reduced starting at 1 DPI and remained so up to 14 DPI. This is at odds with our widespread detection of *Cxcl16* upregulation at 2 DPI and continued upregulation in the CTX at 7 DPI. In brain gliomas, Cxcl16-Cxcr6 receptor signaling in macrophages and microglia promotes an anti-inflammatory cell phenotype [85].

### A subacute transition to adaptive immune cell responses in the contused forebrain

The transcriptional response to injury is downregulated and transitions to include adaptive, B-cell mediated immune responses by 7 DPI. Among the adaptive immune response at 7 DPI are genes involved in immunoglobulin class switching on B cells that suggests B cells encountering antigens in the tissue milieu [86]. This shift in inflammatory and immune responses to injury between an acute 2 DPI and a more subacute 7 DPI timepoint aligns with reviews of the TBI literature demonstrating that by 7 DPI, the injured tissue is undergoing (sometimes failed) repair and experiencing a temporal switch in infiltrating immune cells towards adaptive T/B cells [64, 87, 88] While we see an early upregulation of gene transcripts suggestive of T and B cell infiltration following focal brain injury, we did not measure increases in infiltrating T and B cells nor characterize activation and subset identity of these cells - for example delineating IFN-γ producing Th1 vs anti-inflammatory IL-4 producing Th2 subsets from infiltrating immune cells in the ipsilateral brain hemisphere. However, there is growing support in the literature for T and B cell involvement in acute and chronic TBI pathology.

Il-17 producing CD4^+^ T cells and IFN-γ producing T cells are reported to be elevated by CCI in the contused hemisphere of adult mice at 7 DPI [20], although it should be noted the CCI injury used in this study is more severe than that used in our model (2 mm depth contusion and 5 m/s impact velocity versus 1 mm depth and 1.5 m/s in this study). Interestingly, these authors report depletion of CD8^+^ T cells but not CD4^+^ T cells in CCI mice resulted in an improved neurological outcome and an associated T helper cell type 2 (Th2) response. Furthermore, deletion of B cells in the same study resulted in a heightened immune response in their CCI mice and a progressive deterioration in neurological performance over 32 weeks post-injury.

More recently, early infiltrating gamma-delta (γδ) T cell subsets capable of responding to pattern recognition receptors have been shown to exert subset-specific effects in TBI as early as 1DPI and up to one-year post-injury [19]. Infiltrating V4γδ T cells were shown to secrete IFN-γ and IL-17 to activate microglia and induce neuroinflammation. In contrast, V4γδ T cells secreted TGF-β to maintain microglia homeostasis and suppress neuroinflammation. Again, these authors used a more severe CCI injury than our study, albeit moderately with the same 1 mm depth of contusion but a greater impact velocity of 5 m/s. It will be important to see if these T cell and B cell responses are replicated in less severe, tissue-sparing preclinical models of TBI, but it is noteworthy that *Cd69*, a marker of T cell subsets, including resident memory (TRM) T cells and γδ T cells [89, 90], is prominently upregulated in our injury cluster (cluster 13) at 7 DPI.

### Spatial clustering reveals injury “hotspots” and a layered response to injury in the neocortex

Unbiased spatial clustering of transcriptional responses to CCI revealed features of our data not captured in histologically-defined clusters of the CTX, CC-EC, STR and AC. These features were the presence of an injury cluster or “hotspot” present only in CCI mice at 2 DPI and 7 DPI (clusters 6 and 13 respectively), and a layered, acute transcriptional response to CCI in the CTX (see Figure 4). As expected, injury “hotspots” showed enriched expression of genes related to inflammation, cytokine signaling, and immune cell recruitment. Genes upregulated in the injury hotspot and not in underlying cortical clusters at 2 DPI included the neutrophil chemoattractant *Cxcl1* and the CXC receptors *Ccr2* and *Ccr7*, consistent with early, localized trafficking of neutrophils and monocytes to the injury core. Also, uniquely upregulated in the 2 DPI injury hotspot were the interleukins *Il6*, *Il1* and *Il19* that exert anti-apoptotic, anti-inflammatory effects necessary for re-vascularization and wound healing in the ischemic brain [91–93]. By 7 DPI, there is a clear transition to the localized activation of genes with adaptive immune functions in the injury hotspot (cluster 13). These included the upregulation of *Cxcl9*, a chemoattractant for Ccr3^+^ T lymphocytes and Natural Killer (NK) cells [63], and *Cxcl13*, a chemoattractant for Ccr5^+^ B lymphocytes, the T cell marker Cd69, as well as markers for microglia and macrophage activation such as *Cd5l* and *Siglec1*. Overall, injury hotspot transcriptional signatures support the active transition from innate to adaptive immune cell responses during the first post-injury week in our CCI mice.

Spatial clustering of the laminar CTX revealed a positive relationship between the number of genes differentially expressed within each cluster and the proximity of the cluster to the impact at 2 DPI. This suggests a direct relationship between the initiating physical force and the acute transcriptional response to injury in our CCI mice. Outer cortical layers that experience higher magnitudes of physical force are predicted to sustain greater mechanical disruption of their cellular structures and an increased release of DAMPs. Although the inflammatory effects of DAMPs varies by cell type and the level of expression of pattern recognition receptors [94], spatial differences in the magnitude of DAMP release would be expected to translate to spatial differences in the inflammatory response and in the corresponding magnitude of transcriptional responses in resident CNS cells and in infiltrating immune cells.

### Depletion of immune cell infiltration and metabolic homeostasis in the injured brain

In this study we administered antibodies to granulocyte receptor-1 (GR-1) to systemically deplete circulating neutrophils and monocytes. GR-1 is comprised of two structurally-related glycosylphosphatidylinositol (GPI)-anchored proteins of the Ly-6 superfamily, Ly6G and Ly6C, and the antibody used in this study (clone RB6-8C5) reacts strongly with mouse Ly6G and weakly with mouse Ly6C. This was reflected in flow cytometry data demonstrating reduced infiltration of Ly6G^+^ neutrophils and Ly6C^low-int^ monocytes into the brain at both endpoints but reduced infiltration of Ly6C^hi^ monocytes at only the later 7 DPI endpoint [see **Supplementary Figure 7** data]. Based on these data, we conclude anti-GR-1 treatment preferentially attenuated the acute infiltration of neutrophils and non-classical, patrolling monocytes, thereby skewing early peripheral immune cell responses in the injured brain toward classical, inflammatory Ly6C^hi^ monocytes.

Anti-GR-1 treatment was associated with multifaceted effects on gene pathways, notably an acute upregulation of synaptic signaling and plasticity in the CTX and STR at 2 DPI, and a switch in cytoplasmic translation in mid cortical (cluster 1) layers from downregulation at 2 DPI to upregulation at 7 DPI. Increased transcription of “synaptic” genes in the CTX and STR at 2 DPI is likely a compensatory response to the loss or damage of neural circuits by the CCI [95]. Conversely, the acute downregulation of cluster 1 genes functioning in cytoplasmic translation could be a compensatory reallocation energy of resources: excitotoxicity, oxidative stress, and inflammation induced by a TBI can severely compromise energy production [96] and cytoplasmic translation is one of the most energetically expensive cellular processes.

Divergent anti-GR-1 effects on Ly6C^low-int^ and Ly6C^hi^ monocytes could account for the temporal switch in the relative activity of *cytoplasmic translation* gene pathways with GR-1+ immune cell depletion: downregulated at 2 DPI and upregulated at 7 DPI. Attenuated Ly6G^+^ neutrophil and Ly6C^low-int^ monocyte infiltration at 2 DPI is expected increase post-injury inflammation and oxidative stress relative to IgG treated controls. This could be due to the loss of anti-inflammatory signals from these cells, or the enhanced differentiation of infiltrating Ly6C^hi^ monocytes into pro-inflammatory macrophages in the absence of competing Ly6C^low-int^ monocytes, or from a combination of both factors [97–101]. In this tissue environment, the downregulation of energy intense cytoplasmic translation could be a tradeoff for the energy requirements of the synaptic plasticity of the CTX (and STR). In contrast, attenuation of Ly6C^hi^ monocytes at the later 7 DPI endpoint is expected to reduce inflammation and oxidative stress relative to IgG treated CCI controls, creating a more favorable tissue environment for the post-injury recovery of cytoplasmic translation. Interestingly, increased cytoplasmic translation in cluster 1 at 7 DPI occurred simultaneous to the downregulation of mRNA metabolism, processing and splicing gene pathways at this endpoint. This could reflect the complex interplay between mRNA turnover and translation efficiency designed to optimize protein production under the challenging post-injury environment.

## Conclusion

To conclude, in this study we provide a detailed description of the spatiotemporal characteristics of early transcriptional response to a CCI injury in mice. Our data demonstrates experimental TBI initiates spatial responses in the brain that are shaped by the local cytoarchitecture and proximity to the initiating force of the trauma. In addition, we provide evidence that infiltrating immune cells contribute to the transcriptional regulation of synaptic transmission, metabolism and gene translation post-injury in a spatially-dependent manner. These findings highlight the importance of considering both spatial and temporal factors when developing treatments for TBI.

## Declarations

## Data Availability

Data generated and analyzed in this study will be made available through Figshare, DOI: 10.6084/m9.figshare.27974352

## Ethics declaration

not applicable.

## Disclaimer

The opinions, interpretations, conclusions and recommendation are those of the authors and are not necessarily endorsed by the U.S. Army, Department of Defense, the U.S. Government or the Uniformed Services University of the Health Sciences. The use of trade names does not constitute an official endorsement or approval of the use of reagents or commercial hardware or software. This document may not be cited for purposes of advertisement.

## Author Contributions

SKW contributed to the conceptualization of the study, funding acquisition, the development of study models and methodology, the provision of resources, to data collection, validation and visualization, and to the writing (original draft and review and editing). CA, EG and GS contributed to data acquisition. AF and MDW contributed to data analysis and curation. CLD contributed to resource support, methodology, and data analysis and curation. JTM contributed to resource support. MLD contributed to the conceptualization of the study, funding acquisition, study models and methodology, the provision of resources, project administration and supervision, visualization of the data and to the writing (original draft and review and editing).

## Acknowledgements

We would like to thank Dr. Dennis McDaniel for assistance with imaging and Dr. Kalpana Dommaraju for assistance with bioinformatics data analysis.

## Transparency, Rigor and Reproducibility Statement

The authors affirm that this manuscript is an honest, accurate, and transparent account of the data being reported. In the interests of transparency, rigor and reproducibility, this study provides descriptions of all experimental methods, data analysis pipelines and statistical tests used, in addition to the access of the RNA-sequencing data used in this study to ensure these findings can be readily replicated by other researchers.

## Supplementary Data

**Table A:**
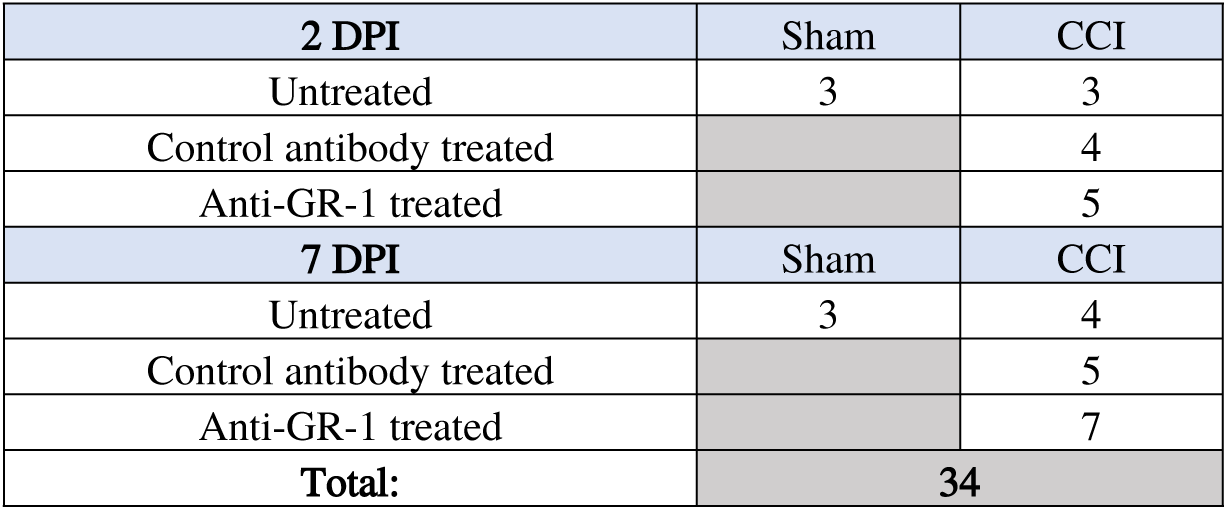
Experimental Groups with Sample Sizes for Visium Spatial Gene Expression.

**Table B:** Antibodies Used for Quantification of GR-1^+^ immune cell depletion by Flow Cytometry.

| Marker | Fluorophore | Clone | Manufacturer | Volume (µL) per Test |
| --- | --- | --- | --- | --- |
| CD45 | Violet Fluor450 | 30-F11 | Cell Signaling Technologies (CST)/Cytek | 2.5 |
| Ly6G | Red Fluor 710 | 1A8 | CST/Cytek | 1.25 |
| Ly6C | FITC | HK1.4 | CST/Cytek | 0.25 |
| CD11b | BUV805 | M1/70 | BD Biosciences | 0.63 |
| SiglecF | APC-Cy7 | E50-2440 | BD Biosciences | 1.25 |
| CD182 (CXCR2) | APC | V48-2310 | BD Biosciences | 5.0 |
| CD200R3 | PerCP/Cy5.5 | Ba13 | BioLegend | 0.63 |
| CD11c | PE/Cy7 | N418 | BioLegend | 1.25 |
| CD192 (CCR2) | BV605 | SA203G11 | BioLegend | 3.7 |
| CD3 | BUV563 | 145-2C1 | BD Biosciences | 5.0 |
| CD19 | Spark NIR 685 | 6D5 | BioLegend | 0.5 |
| CD86 | PE/Cy5 | GL1 | Invitrogen | 0.075 |
| CX3CR1 | BV785 | SA011F11 | BioLegend | 0.63 |
| Brilliant Stain Buffer |  |  | BD Biosciences | 50.0 |

**Supplementary Figure 1.**
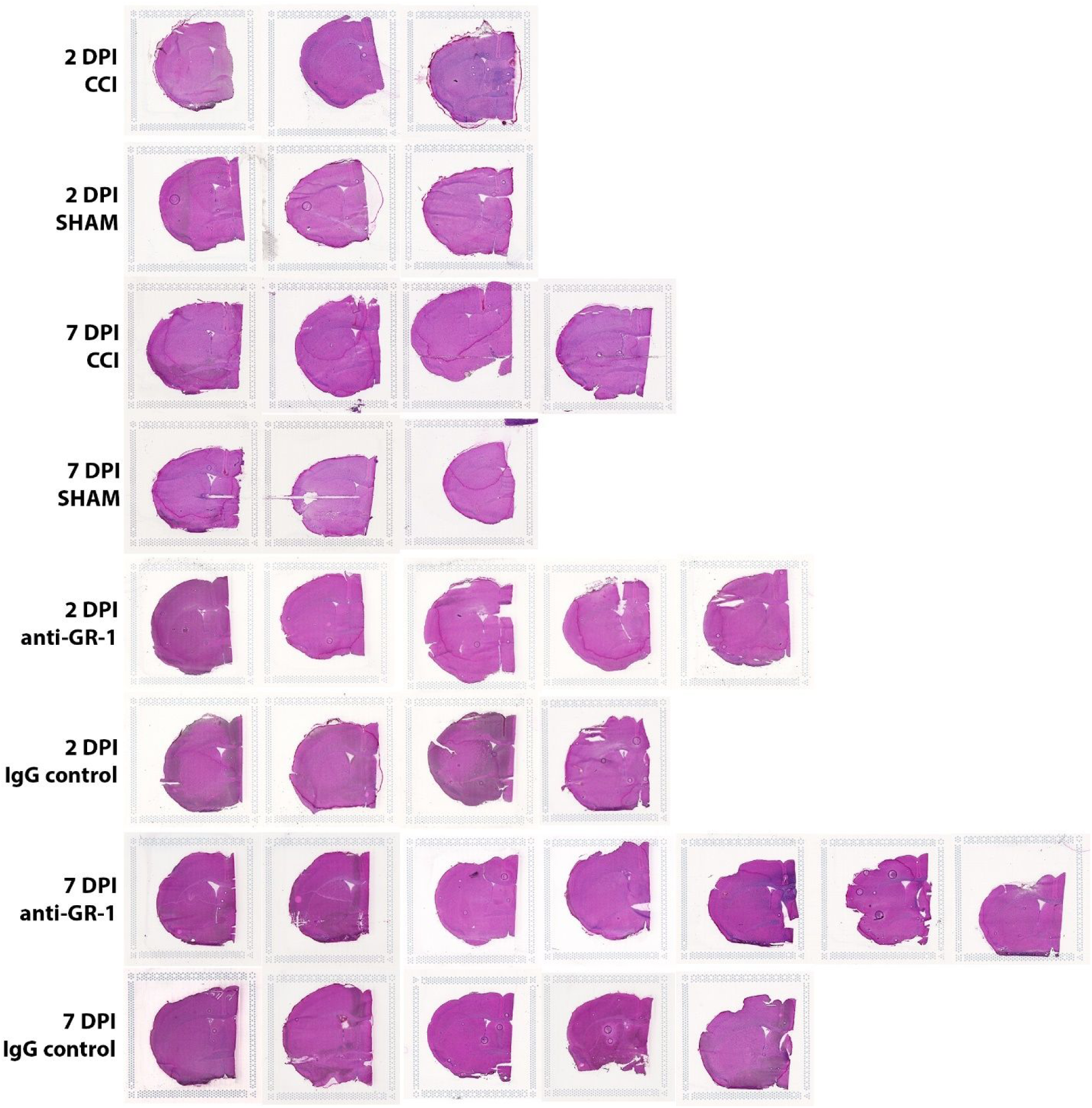
Images of hematoxylin and eosin (H&E) stained coronal sections of the left forebrain of mice used in this study. Coronal sections (12 um thick) were collected at the rostro-caudal level of the anterior commissure for H&E staining, imaging and Visium spatial gene expression analysis.

**Supplementary Figure 2.**
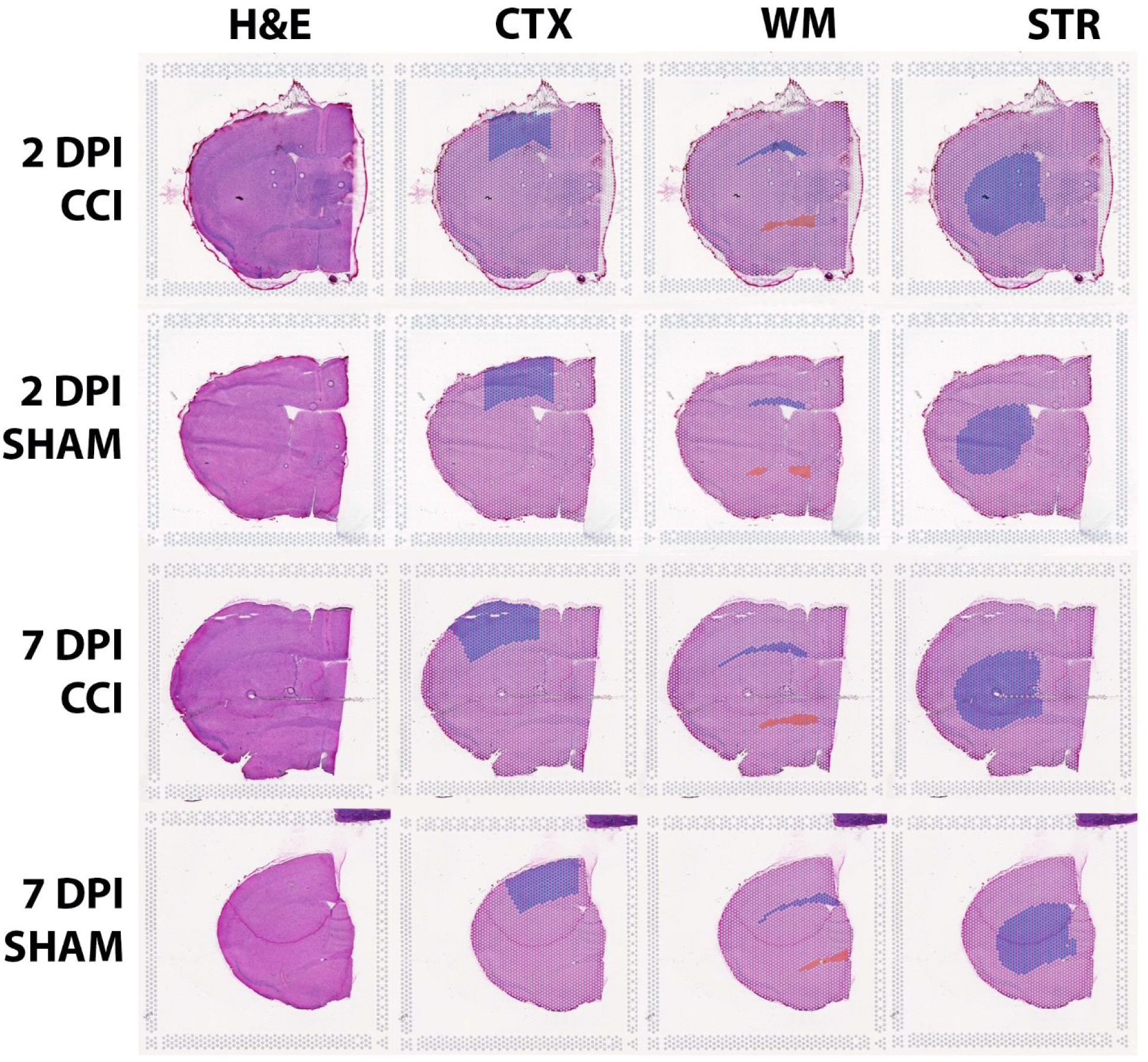
Loupe browser images of (H&E) stained coronal sections and the cortex, white matter and striatum territories used for pseudobulk gene expression comparisons between CCI and Sham mice. **Loupe browser images of H&E stained coronal forebrain sections used for** Visium spatial gene expression analysis. Adjacent are images showing the annotated cortex, white matter (blue corpus callosum-external capsule, red anterior commissure) and striatum delineated for pseudobulk gene expression comparisons between and within groups.

**Supplementary Figure 3.**
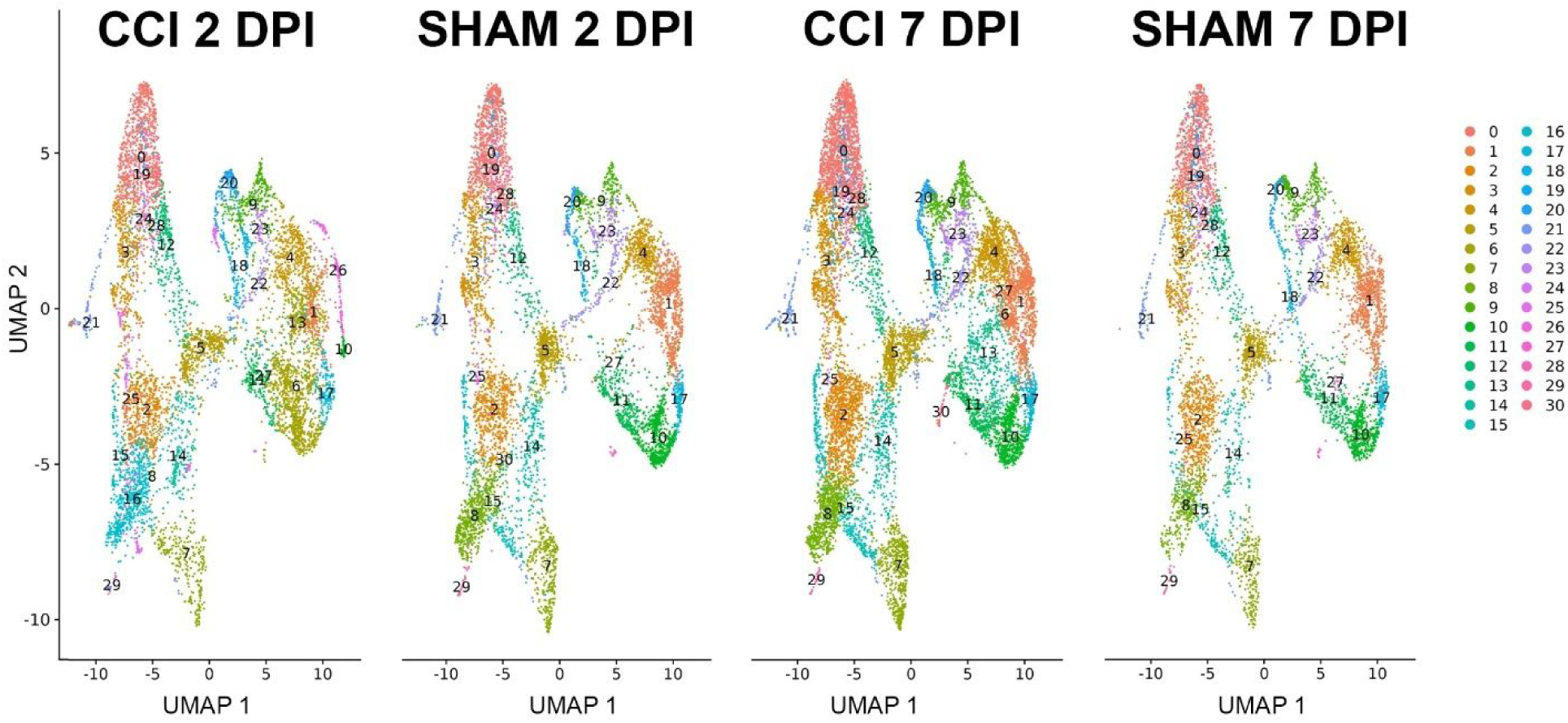
Uniform manifold approximation and projection (UMAP) visualization of spatial clusters demonstrates the gene expression from CCI and sham mice clusters into 29 groups.

**Supplementary Figure 4.**
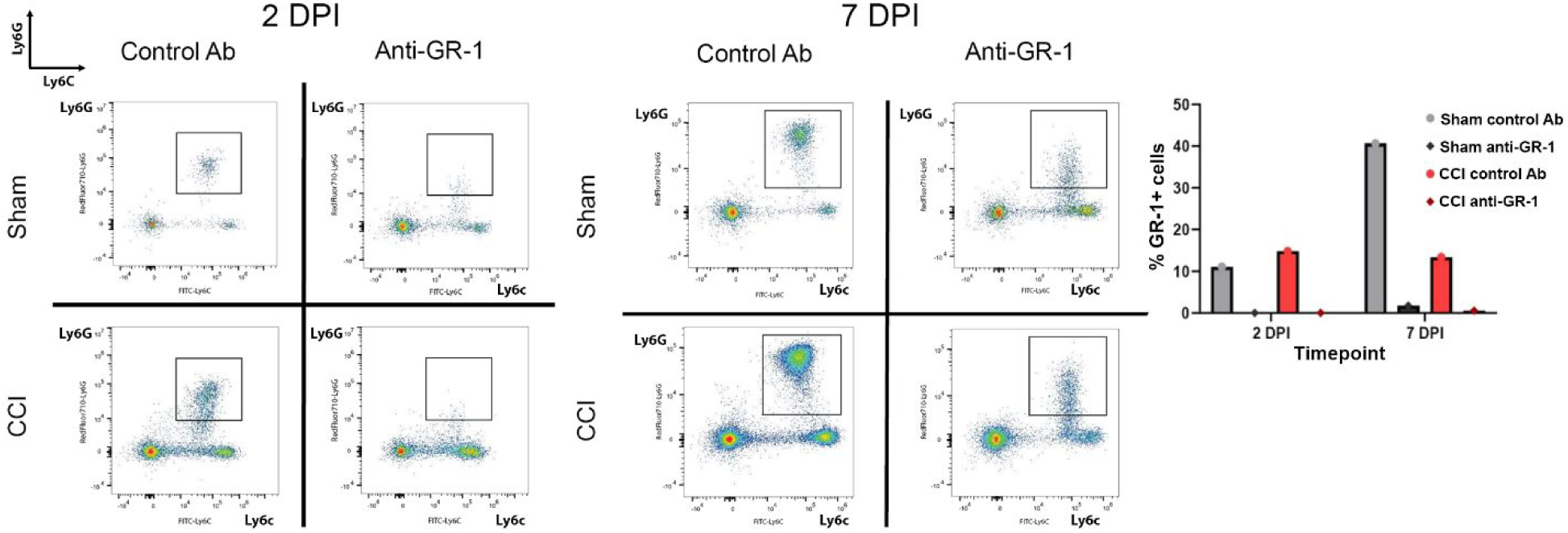
Antibody-mediated depletion of GR-1^+^ cells systemically depletes circulating leukocytes in CCI and sham mice. Mice were treated by with 250 µg per day of anti-GR-1 antibody or the same amount of an isotype control antibody by intraperitoneal injection beginning 1 day prior to CCI or sham intervention and continuing every 24 hours until the 2 or 7 DPI endpoint. For validation of GR-1^+^ cell depletion, leukocytes were isolated from blood and spleen at 2 and 7 DPI (data shown for spleen only). For all flow cytometry plots, the images demonstrate cells with x axis = Ly6C positivity in the FITC channel and the y axis = Ly6G positivity in the RedFluor710 channel. For all flow cytometry analyses, n = 2-4 pooled tissues from individual mice in a group for a single run. Pseudocolor plots with Ly6C^hi^/ Ly6G^hi^ populations boxed. Histogram is the percentage of GR-1^+^ cells as calculated from the number of Ly6G^+^/Ly6C^+^ cells divided by the number of live CD45^+^ cells in CCI or Sham mice treated with anti-GR-1 or control antibody at 2 and 7 DPI.

**Supplementary Figure 5.**
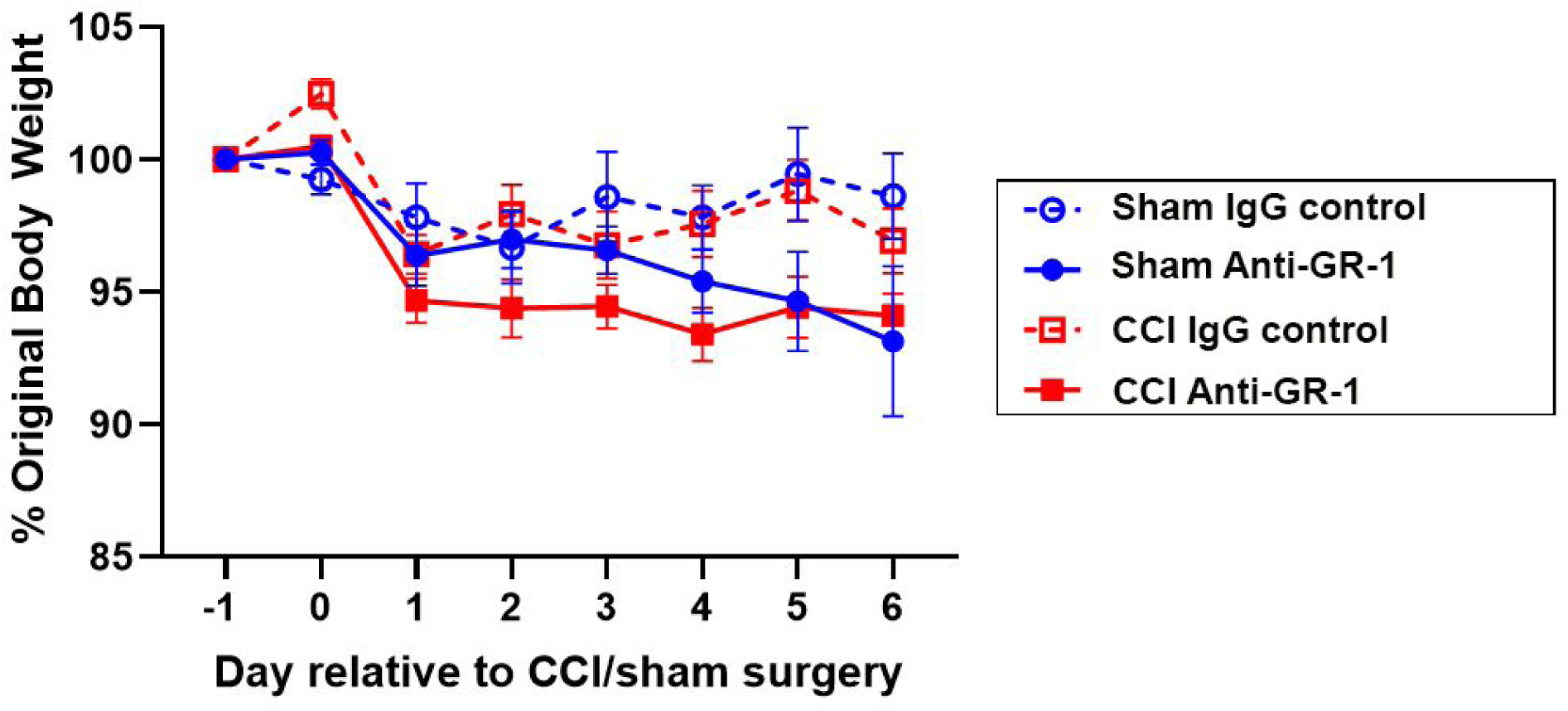
Systemic administration of anti-GR-1 antibodies results in a small but significant loss of weight in CCI and sham mice. Mice receiving anti-GR-1 or control antibody treatment were weighed daily beginning just prior to ip injection until the day before reaching their study endpoint. Anti-GR-1 administration resulted in a small but significant reduction in body weight (mixed effects ANOVA, p < 0.01). Data are mean ± SEM.

**Supplementary Figure 6.**
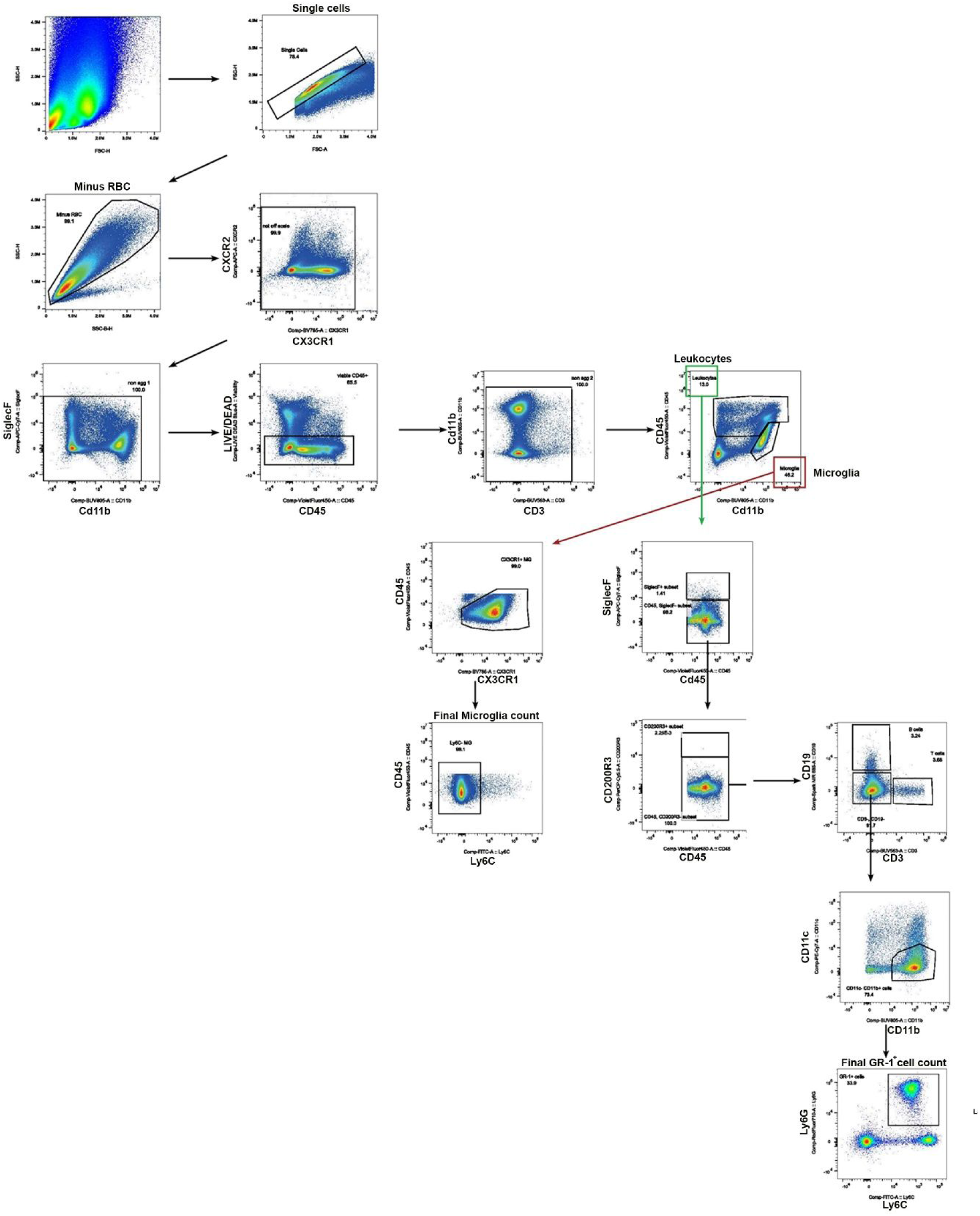
Flow gating strategy to identify microglia and infiltrating peripheral immune cells in the ipsilateral contused brain hemisphere. The microglia population was gated on CD45^low-int^/CX3CR1^+^/Ly6C^−^. Infiltrating leukocytes were first gated on CD45^hi^. Eosinophils and basophils were gated out using SiglecF^+^ and CD200R3^+^ respectively before selecting for the CD3^−^/CD19^−^ population, and then the CD11b^+^/CD11c^−^ cells were selected within this population. GR-1^+^ cells were gated on Ly6G^+^/Ly6C^+^.

**Supplementary Figure 7.**
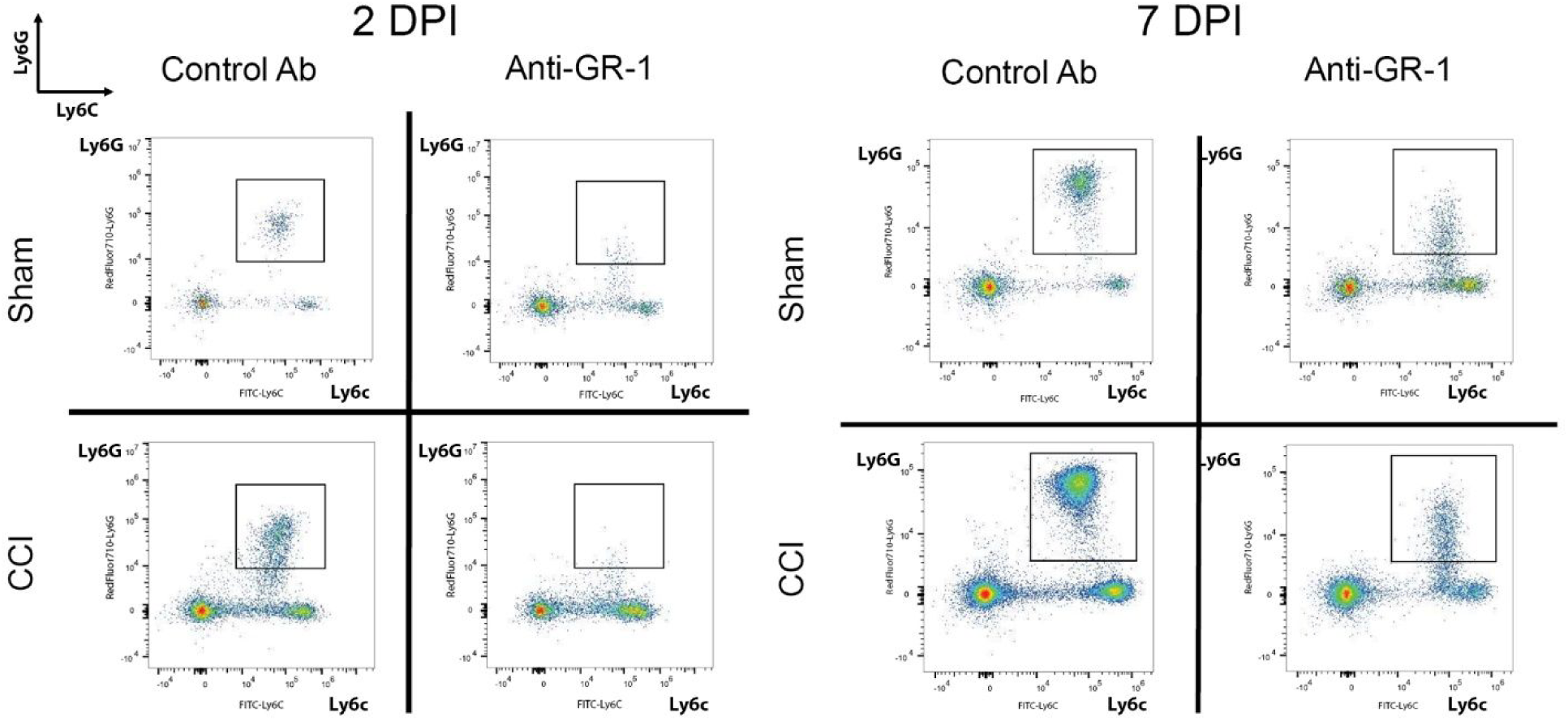
Antibody-mediated depletion of circulating GR-1^+^ cells attenuates the infiltration of Ly6G^+^/Ly6C^+^ leukocytes into the damaged brain post-injury. Pseudocolor plots reveal robust depletion of Ly6G^hi^/Ly6C^hi^ leukocytes (black boxes) at 2 and 7 DPI. Anti-GR-1 treatment reduced the number of infiltrating Ly6G^+^ neutrophils and Ly6C^low-int^ monocytes at 2 and 7 DPI, and the infiltration of Ly6C^hi^ monocytes at 7DPI but not at the earlier 2DPI endpoint.

**Supplementary Figure 8.**
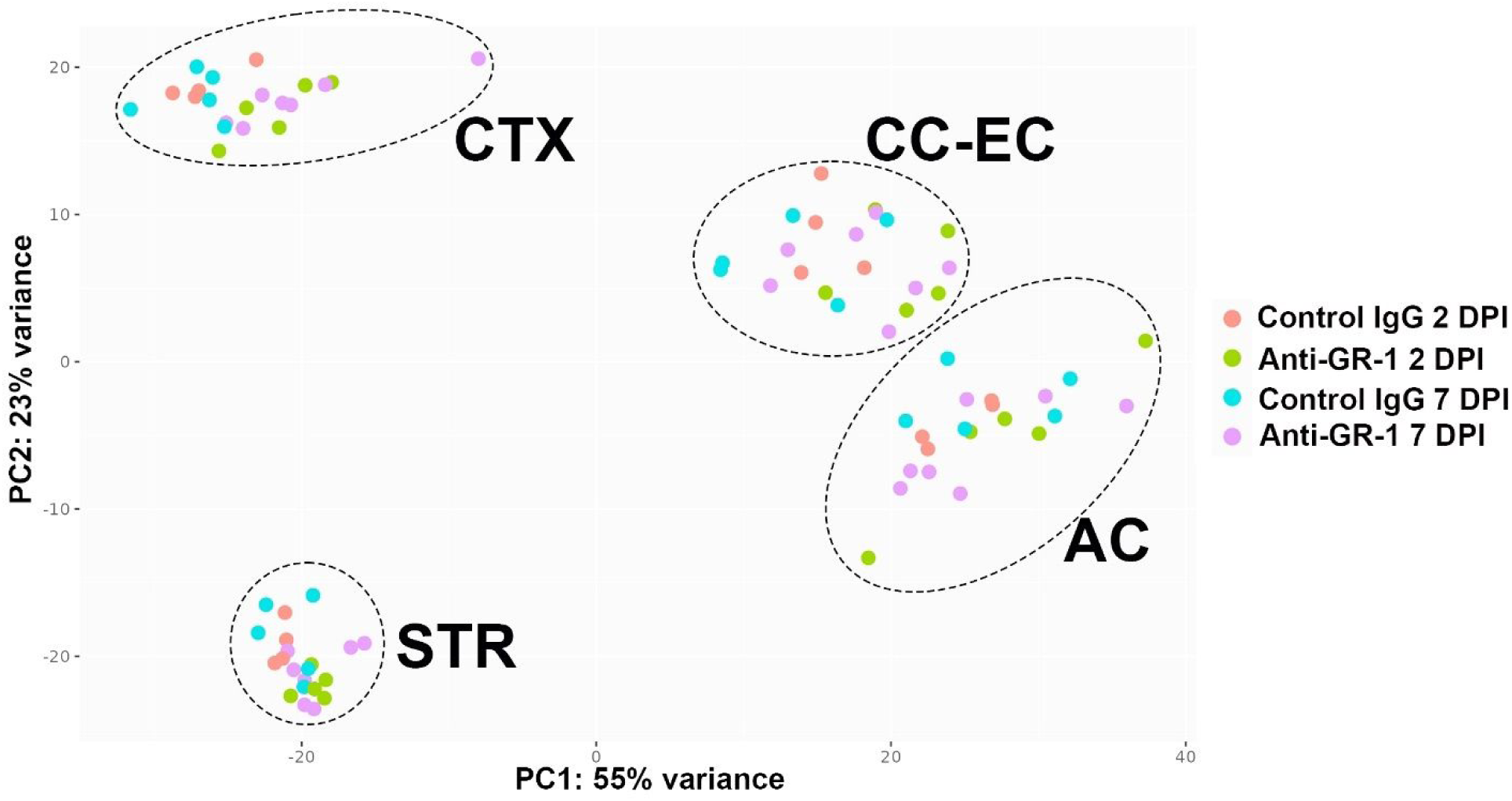
Principal component analysis (PCA) separates pseudobulk DEG data by anatomical territory but not by antibody treatment.

**Supplementary Figure 9.**
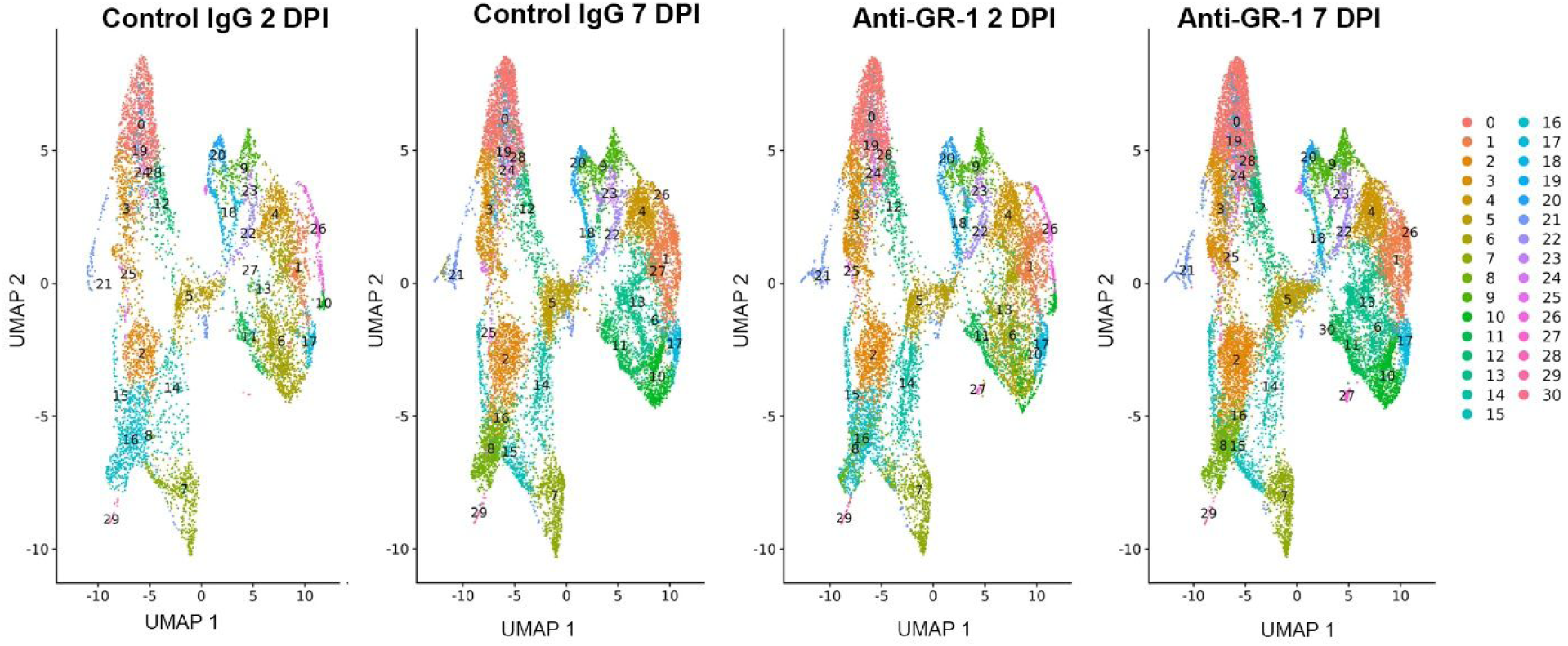
Uniform manifold approximation and projection (UMAP) reveals 29 distinct spatial clusters in the Anti-GR-1 and Control IgG treated CCI mice.

---

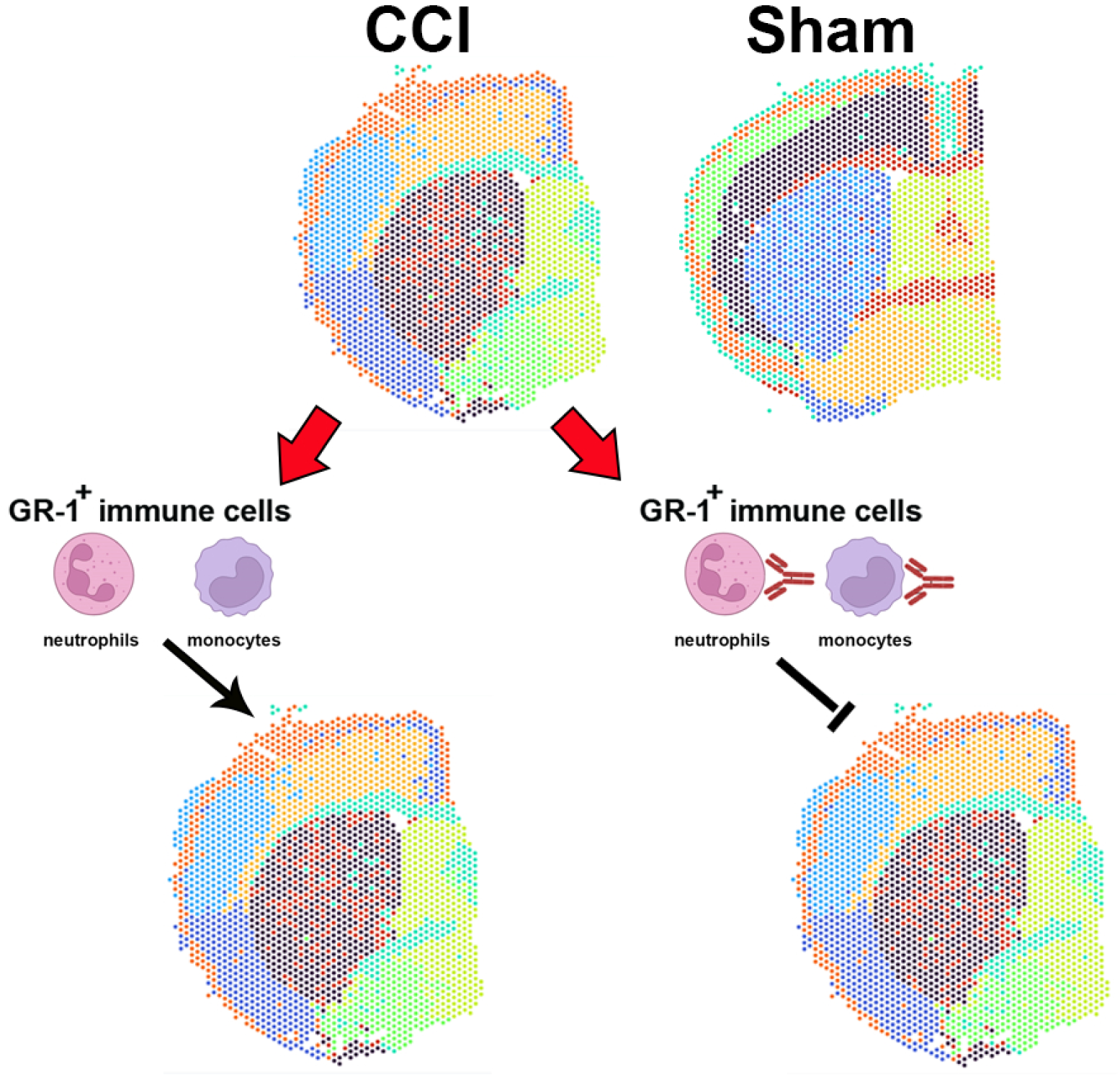

